# Sm-like protein Rof inhibits transcription termination factor ρ by binding site obstruction and conformational insulation

**DOI:** 10.1101/2023.08.30.555460

**Authors:** Nelly Said, Mark Finazzo, Tarek Hilal, Bing Wang, Tim Luca Selinger, Daniela Gjorgjevikj, Irina Artsimovitch, Markus C. Wahl

**Affiliations:** Freie Universität Berlin, Institute of Chemistry and Biochemistry, Laboratory of Structural Biochemistry, Takustr. 6, D-14195 Berlin, Germany; The Ohio State University, Department of Microbiology and Center for RNA Biology, Columbus, OH, USA; Freie Universität Berlin, Institute of Chemistry and Biochemistry, Research Center of Electron Microscopy and Core Facility BioSupraMol, Fabeckstr. 36a, 14195 Berlin, Germany; Helmholtz-Zentrum Berlin für Materialien und Energie, Macromolecular Crystallography, Albert-Einstein-Str. 15, D-12489 Berlin, Germany

## Abstract

Transcription termination factor ρ is a hexameric, RNA-dependent NTPase that can adopt active closed-ring and inactive open-ring conformations. The Sm-like protein Rof, a homolog of the RNA chaperone Hfq, inhibits ρ-dependent termination *in vivo* but recapitulation of this activity *in vitro* has proven difficult and the precise mode of Rof action is presently unknown. Our electron microscopic structures of ρ-Rof and ρ-RNA complexes show that Rof undergoes pronounced conformational changes to bind ρ at the protomer interfaces, undercutting ρ conformational dynamics associated with ring closure and occluding extended primary RNA-binding sites that are also part of interfaces between ρ and RNA polymerase. Consistently, Rof impedes ρ ring closure, ρ-RNA interactions, and ρ association with transcription elongation complexes. Structure-guided mutagenesis coupled with functional assays confirmed that the observed ρ-Rof interface is required for Rof-mediated inhibition of cell growth and ρ-termination *in vitro*. Bioinformatic analyses revealed that Rof is restricted to Pseudomonadota and that the ρ-Rof interface is conserved. Genomic contexts of *rof* differ between *Enterobacteriaceae* and *Vibrionaceae,* suggesting distinct modes of Rof regulation. We hypothesize that Rof and other cellular anti-terminators silence ρ under diverse, but yet to be identified, stress conditions when unrestrained transcription termination by ρ would be lethal.

## Introduction

Bacterial transcription termination factor, ρ, is a hexameric, NTP-dependent 5’-to-3’ RNA translocase and helicase. A ρ monomer contains N-terminal (NTD) and C-terminal (CTD) domains connected by a flexible linker (1). The NTD comprises an N-terminal three-helix bundle and a five-stranded β-barrel, the OB-fold, that forms a primary RNA-binding site (PBS); the PBSes are located around one outer rim of the ρ hexamer. The CTDs jointly form a secondary RNA-binding site (SBS) at the center of the ρ hexamer. The ligand-free ρ hexamer adopts an inactive, open-ring conformation, in which it can bind single-stranded RNA (or DNA) at the PBSes (2). When NTP (preferably ATP) binds to pockets located between the ρ subunits and RNA binds to the SBS, the ring closes, trapping RNA in the central pore of the hexamer and activating the ρ ATPase (3–5). Binding of pyrimidine-rich nucleic acids to the PBSes promotes the formation of a closed-ring active state (6).

ρ triggers premature release of antisense, horizontally-acquired, and untranslated RNAs from transcription complexes, ensuring that useless or potentially harmful RNAs are not transcribed (1). Productive RNA synthesis requires protection of transcription complexes from ρ. When RNA and protein synthesis are coupled, such as in fast-growing *Escherichia coli*, a translating lead ribosome is the main line of defense against ρ, as it denies ρ access to the transcribing RNA polymerase (RNAP) or nascent RNA (7). Bacterial RNAs that are translated poorly (e.g., transcripts of xenogeneic cell wall biosynthesis operons) or not at all (e.g., ribosomal [r] RNAs) require dedicated anti-termination mechanisms to avoid being terminated by ρ (1). Bacteriophages, pervasive mobile elements that are silenced by ρ (8, 9), use analogous mechanisms to ensure that their genes are expressed. Anti-termination is mediated by large ribonucleoprotein complexes, *e.g.*, during rRNA synthesis or phage λ gene expression (10, 11), single proteins such as RfaH (12, 13), or RNAs (14–16), which are recruited to specific sequences or encoded within their target operons and inhibit termination within just one or a few operons.

It is currently unknown how ρ activity is regulated during periods of slow growth or stress, when translation is inefficient and uncontrolled transcription termination may be lethal. Cellular levels of ρ are kept almost constant (17) through auto-regulation (18), implying that during stress, termination may be kept in check by regulators that reduce ρ activity. Three cellular anti- termination factors have been reported to inhibit ρ; the RNA chaperone Hfq, the Ser/Thr kinase YihE, and the small protein YaeO (a.k.a. Rof in *E. coli*). Hfq can directly bind ρ, and inhibits ρ by an unknown, and possibly indirect, mechanism (16, 19). YihE forms a complex with ρ and inhibits ρ-RNA interactions (20). Rof proteins adopt an Sm-like fold, resembling Hfq (Fig. 1*A*), and also directly bind ρ (21, 22). While NMR-based docking and RNA binding studies suggested that Rof also competes with RNA for binding to ρ (21), the precise mode of the ρ-Rof interaction and of Rof-mediated ρ inhibition remain unclear.

**Figure 1.**
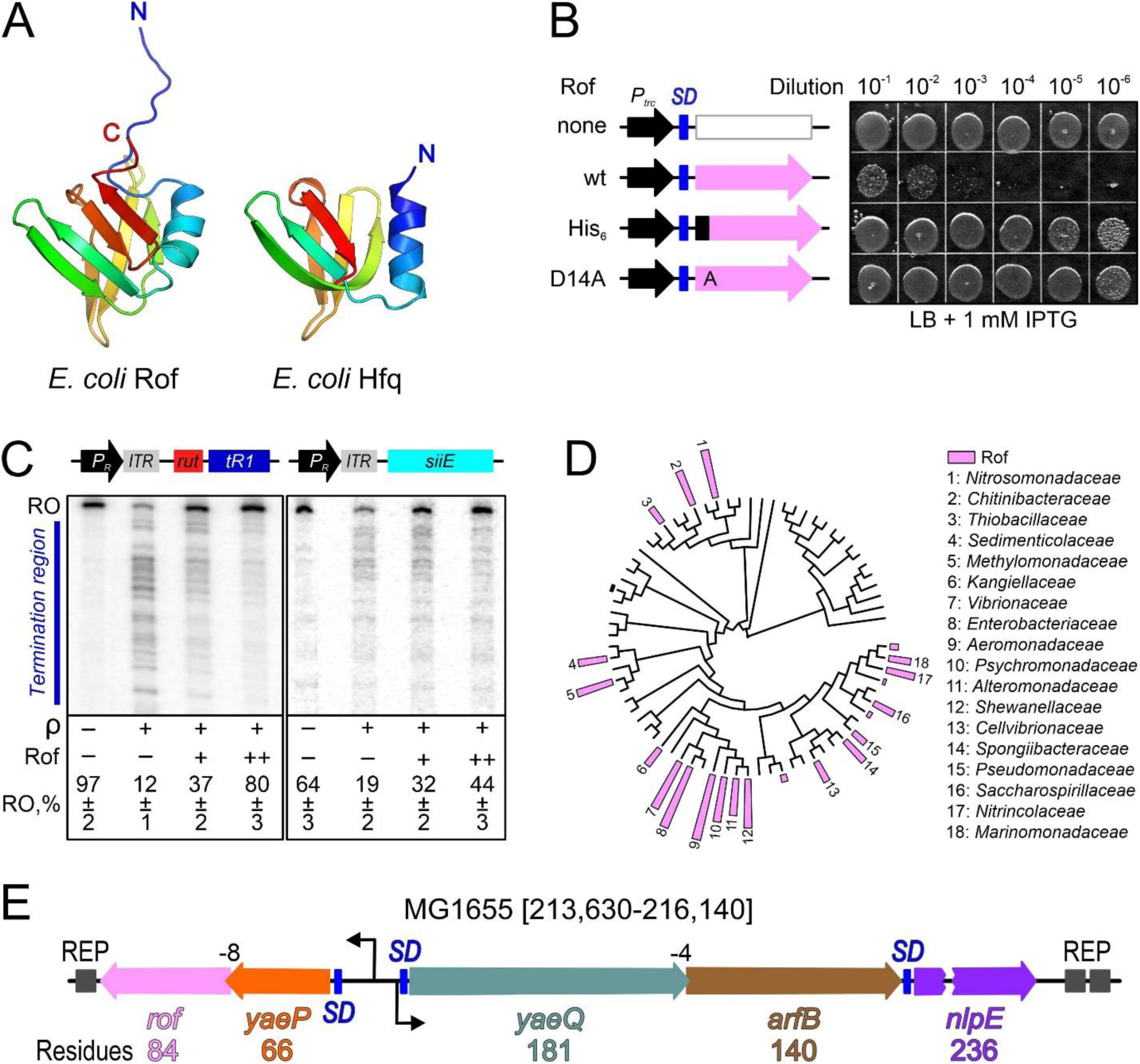
(A) Structural comparison of *E. coli* Rof (left) and Hfq (right). Proteins are shown in rainbow colors from N-termini (N; blue) to C-termini (C; red). (B) Cells transformed with plasmids expressing wildtype (wt) Rof and Rof variants under the control of an IPTG-inducible P*_trc_* promoter or an empty vector were grown overnight in LB supplemented with carbenicillin at 32 °C. Serial 10-fold dilutions were plated on LB-carbenicillin in the presence of IPTG and incubated overnight at 32 °C. A set representative of five independent experiments is shown. Shine-Dalgarno (*SD*) elements are indicated by blue bars. (C) Rof inhibits ρ-dependent termination *in vitro*. Top, DNA templates encoding the *rut*^+^ λ *tR1* (left) or *rut^-^ siiE* leader region (right) downstream of the λ *P_R_* promoter and a C-less cassette that allows for the formation of synchronized, radiolabeled elongation complexes halted 26 nts downstream of the transcription start site (arrow). Bottom, representative single-round transcription reactions. Halted elongation complexes were incubated with ρ and/or Rof (where indicated), restarted by addition of NTPs, and subsequently quenched. Reactions were analyzed on 5 % urea-acrylamide gels. Positions of the full-length run-off (RO) RNA and termination regions are indicated. The fractions of run-off RNA represent the mean ± SD of three independent experiments. (D) Rof distribution on the phylogenetic tree of Pseudomonadota. Each leaf represents one family. The height of the bar indicates the fraction of Rof in each family, with the highest one being 80 %. (E) *E. coli yae* locus (MG1655 genome coordinates 213,630-216,140); the length of each ORF is indicated below the sequence, the *nlpE* gene is interrupted to save space. The *arfB* and *rof* genes lack Shine-Dalgarno elements and overlap with the respective upstream ORFs, *yaeP* and *yaeQ*, by 4 and 8 nts. The “insulating” REP elements are shown by dark gray boxes.

Here, we used a combination of structural, biochemical, and genetic approaches to delineate in detail the mechanism of Rof-mediated inhibition of ρ. We show that Rof reduces ρ-dependent termination *in vitro* and inhibits *E. coli* growth, presumably by interfering with the essential functions of ρ. We present cryogenic electron microscopy (cryoEM)/single-particle analysis (SPA)-based structures of ρ-Rof complexes, precisely defining the ρ-Rof interfaces and showing that Rof undergoes substantial conformational changes upon binding to ρ. Substitutions of interface residues interfere with Rof activity *in vitro* and *in vivo,* confirming that the observed molecular interactions are required for ρ inhibition. Rof binds near the canonical ρ PBSes, but does not seem to directly compete with the binding short oligonucleotides (2). To further explore the mechanism of Rof-mediated inhibition of ρ, we obtained a structure of ρ bound to a 99-nt long RNA that occupies all RNA-binding sites on ρ. This structure revealed a previously proposed extended PBS that is blocked upon Rof binding. Furthermore, structural comparisons showed that Rof undercuts functional communication between ρ NTDs and CTDs *via* the connecting linker. In addition, we show that Rof prevents ρ binding to a transcription elongation complex (EC). Thus, Rof blocks termination by inhibiting ρ recruitment to either the naked RNA or to ECs. Finally, our bioinformatics analyses of Rof distribution across bacteria and its genomic locations suggest that Rof and other known or yet to be identified anti-terminators comprise a set of regulators that mediate stress-specific responses by attenuating ρ activity.

## Results

### Rof is toxic *in vivo* and prevents ρ-dependent transcription termination *in vitro*

A pioneering study of *E. coli* Rof had shown that, when overexpressed, Rof co-purifies with ρ and induces readthrough of a canonical ρ terminator in the cell; surprisingly, however, purified Rof did not inhibit termination *in vitro* (23). We wondered if the presence of an N-terminal His-tag could have been responsible for this apparent loss of inhibition. The cellular function of Rof is not yet known, prompting us to establish a proxy assay for Rof activity *in vivo*. We previously showed that ectopic expression of Psu, an unrelated phage P4 inhibitor of ρ, directed by an IPTG-inducible P*_trc_* promoter was toxic (24); we, thus, constructed a plasmid with the *rof* open reading frame (ORF) under the control of P*_trc_*. A plasmid carrying wildtype (wt) *rof*, but not an empty vector, strongly inhibited growth of IA227 (25), a derivative of MG1655 (Fig. 1*B*). Rof-mediated toxicity was abolished by the addition of an N-terminal His-tag or substitution of D14, a conserved Rof residue shown to be critical for the ρ-Rof interaction (21), for alanine (Fig. 1*B*).

The above results suggest that Rof inhibits the essential functions of ρ. One of the critical roles of ρ is to silence prophages, and MDS42, a genome-reduced *E. coli* strain from which all cryptic prophages have been removed by targeted deletion (26), is ∼10^4^ times more resistant to bicyclomycin, which counteracts ρ ring closure (6), than the parent MG1655 strain (8). We found that MDS42 was highly resistant to Rof-mediated killing (Fig. S1*A*), and Morita *et al*. reported that Rof and bicyclomycin have similar effects on synthesis of a small RNA sgrS (27). Together, these result support a model that Rof acts as a general inhibitor of ρ and that Rof toxicity is alleviated under conditions that reduce the cell dependency on ρ.

To determine at what stage of cellular growth Rof expression is inhibitory, we monitored the growth kinetics of IA227. We found that Rof^wt^ extended the lag phase from 2.75 to 12.65 hours but did not alter the maximal growth rate or the saturation point (Fig. S1*B*). The extended lag could reflect defects in protein synthesis required to sustain rapid growth (28, 29) or loss of viability of the seed culture in the stationary phase. In support of the second explanation, we found that Rof expression dramatically reduced viable colony count (Fig. 1*B* and S1*A*). As Rof expression inhibited the growth of MDS42 only modestly (∼1-log; Fig. S1*A*), we conjecture that the observed loss of IA227 viability could be due to the activation of cryptic prophages, a scenario that we are actively investigating.

While our *in vivo* results (Fig. 1*B*) support the hypothesis that the failure to observe anti-termination *in vitro* (23) could be due to the presence of the His-tag on Rof, it is also possible that Rof, proposed to compete with RNA for binding to ρ (21), may not be able to outcompete a tightly bound RNA; the failed *in vitro* trials (23) used λ *tR1*, one of the highest-affinity ρ ligands known. To test this idea, we compared effects of tag-less Rof *in vitro* on templates carrying λ *tR1* and a leader region of *Salmonella siiE,* which is inhibited by ρ *in vivo* but lacks C-rich *rut* elements preferentially recognized by ρ (30). We found that ρ terminated RNA synthesis on both templates (Fig. 1*C*), supporting a view that the *rut* site is less important in the context of the transcription complex (31, 32). Tag-less Rof efficiently counteracted ρ-dependent termination on both templates, suggesting that an N-terminal His-tag indeed interferes with Rof activity and arguing that the Rof effects on ρ are direct and independent of *rut* sites (Fig. 1*C*).

### Rof binds to ρ *via* a conserved interface

Consistent with pull-down assays (23), we found that Rof forms a stable complex with ρ in analytical size-exclusion chromatography (SEC), as a mixture of both proteins eluted at an earlier volume than either protein alone (Fig. 2*A*). Multi-angle light scattering (MALS) detection revealed that ρ and the ρ-Rof complex had heterogeneous molecular mass distributions (Fig. 2*A*), consistent with earlier observations that ρ does not exist as a discrete hexamer in solution (33).

**Figure 2.**
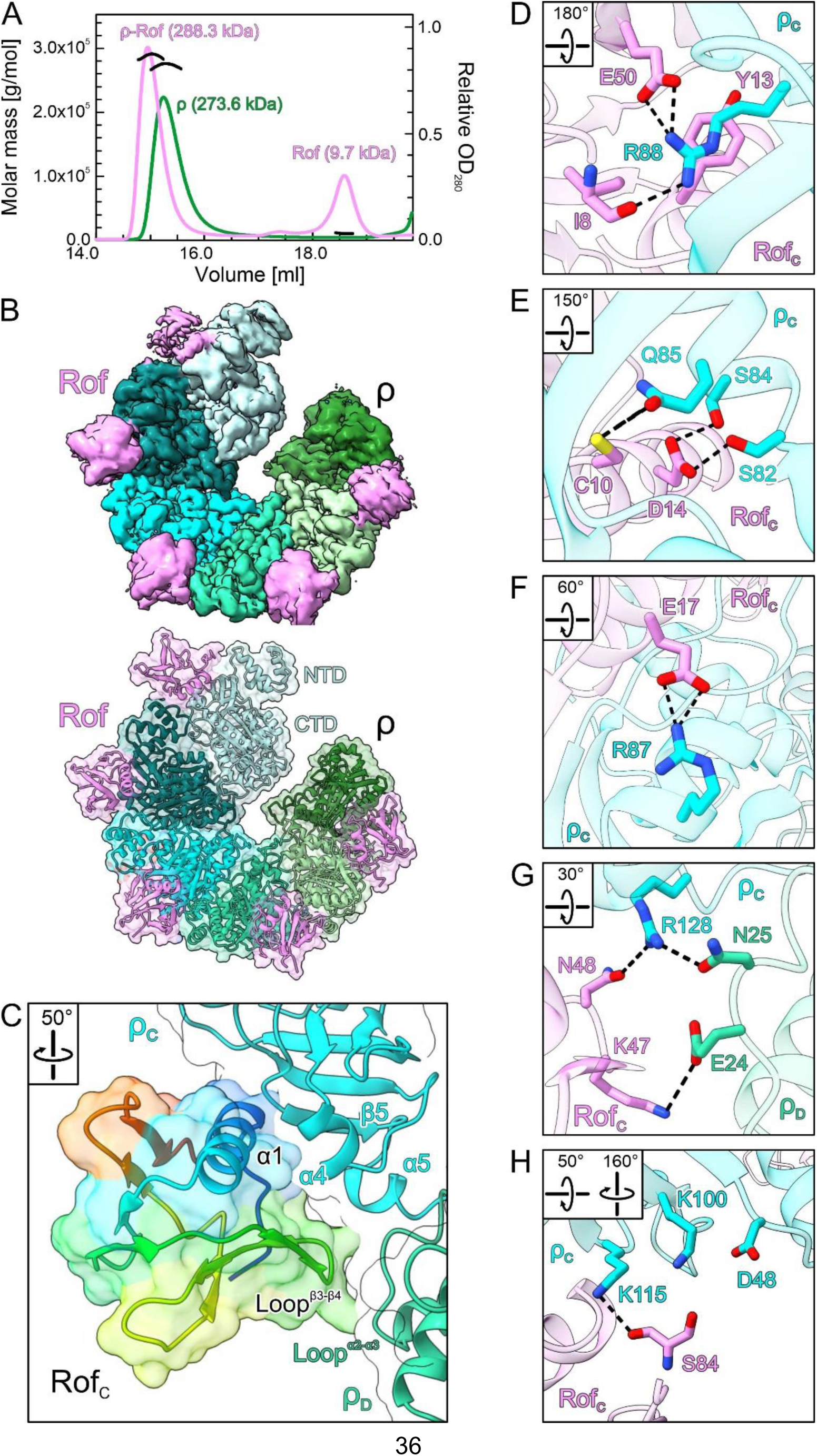
(A) SEC-MALS of isolated ρ (green) and ρ in the presence of Rof (violet). The average molar masses of the peak fractions are indicated. (B) CryoEM reconstruction (top) and cartoon/surface representation (bottom) of the ρ_6_-ADP-Rof_5_ complex. ρ, different shades of cyan and green; Rof, violet. ρ adopts an open ring conformation and Rof proteins bind between the NTDs of neighboring ρ protomers. (C) Close-up view of the ρ-Rof interface. Rotation symbols in this and (*D-H*) indicate the views relative to (*B*). Rof is shown in rainbow colors from N-terminus (blue) to C-terminus (red). Rof binds ρ *via* its N-terminal helix α_1_, the β3-β4 loop and the C-terminus. (*D-H*) Details of the ρ-Rof interaction. Interacting residues are shown as sticks with atoms colored by type; carbon, as the respective protein subunit; nitrogen, blue; oxygen, red. Black dashed lines, hydrogen bonds or salt bridges.

We subjected SEC fractions corresponding to ρ-Rof complexes to cryoEM/SPA in the absence and presence of the transition state analog, ADP-BeF_3_ (Fig. S2-S5). Multi-particle 3D refinement with ∼ 600,000 particle images yielded two cryoEM reconstructions for each sample (Fig. S3 and S5), corresponding to two ρ-Rof assemblies. In the first assembly, ρ forms an open hexameric ring bound by five Rof molecules, while the second assembly represents a ρ pentamer bound by four Rof molecules (Fig. 2*B*). In the presence of ADP-BeF_3_, we obtained a ∼ 1:1 (ρ_6_-Rof_5_:ρ_5_-Rof_4_) distribution between the two classes, whereas in the absence of the nucleotide, the equilibrium was shifted towards the lower oligomeric assembly (ρ_6_-Rof_5_:ρ_5_-Rof_4_ ∼ 1:3). For nucleotide-bound complexes, only the ADP portion was visible in the densities. As ρ-Rof interactions are identical in the two assemblies irrespective of bound nucleotide, we focus our subsequent descriptions on the ADP-bound ρ_6_-Rof_5_ complex.

**Figure 3.**
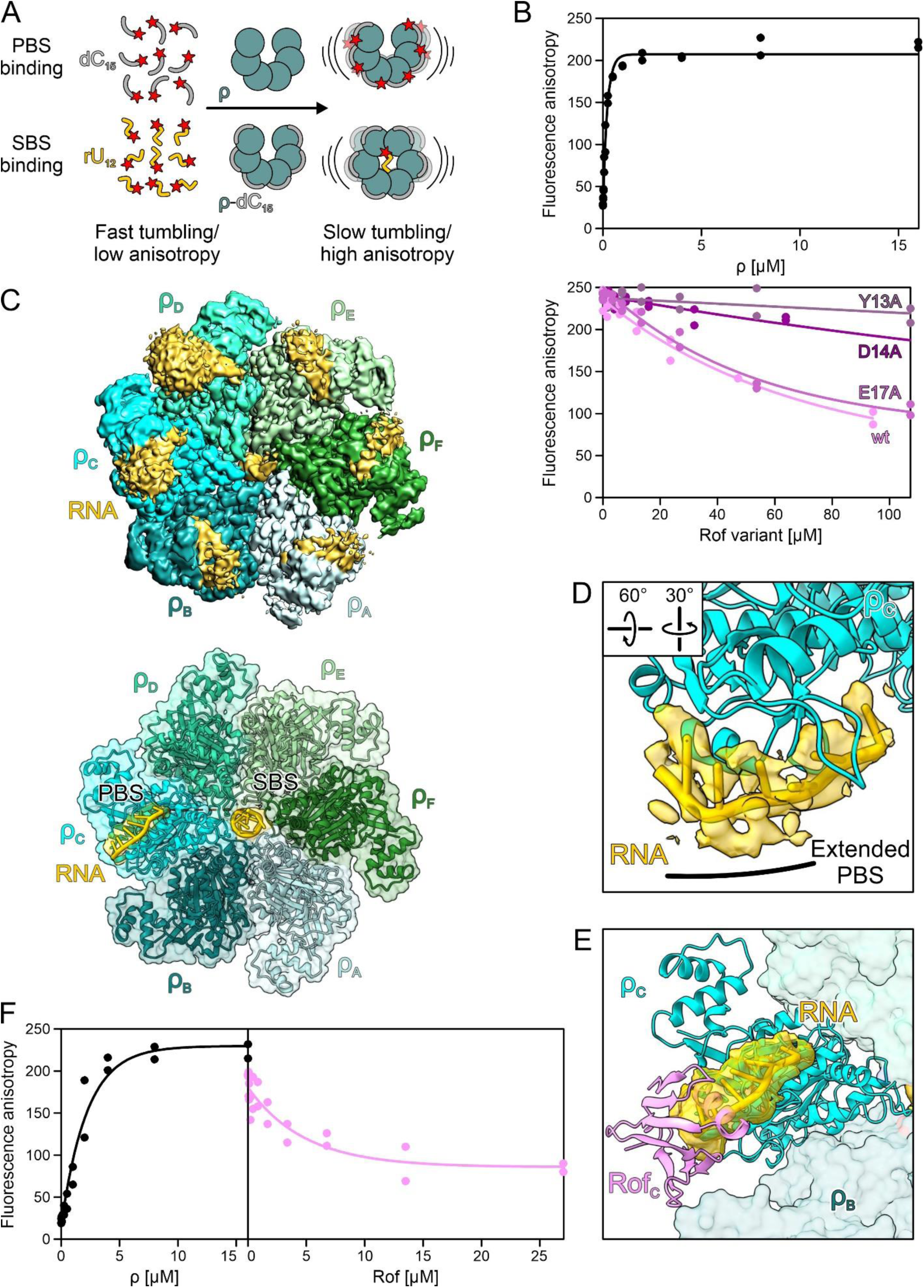
(A) Principles of fluorescence anisotropy assays used in (*B*) and (*G*). Isolated fluorescence-labeled (red star) dC_15_ DNA oligos (gray) or rU_12_ RNA oligos (gold) exhibit high tumbling rates (low fluorescence anisotropy). In the presence of ρ, dC_15_ binds to the PBSes of ρ, which reduces its tumbling rate (increased fluorescence anisotropy). In the presence of dC_15_, rU_12_ binds to the SBS of ρ, which leads to ρ ring closure. Scheme adapted from (64). (B) PBS binding. Top, binding of dC_15_ to the ρ PBSes in the presence of increasing concentrations of ρ. Bottom, effect of Rof variants on the binding of dC_15_ to the PBSes. Data of two independent experiments and fits to a single-exponential Hill function (see Materials and Methods) are shown. (C) CryoEM density (top) and cartoon/surface representation (bottom) of the ρ-*rut* RNA structure. ρ forms a closed ring and is bound by RNA at the NTD of each protomer and at the SBS. Only RNA regions bound at the SBS and at the ρ_c_ NTD were modeled into the cryoEM reconstruction. RNA density and model, gold. (D) ρ interaction with RNA at the PBS. RNA in cartoon and stick representation modeled into the density observed at ρ_c_. The bound RNA 5’-region outlines an extended PBS. Rotation symbols, view relative to (*C*). (E) Structural comparison between ρ-*rut* RNA and ρ_6_-ADP-Rof_5_ complexes. Structures of the complexes were superimposed based on the ρ_c_ subunits. Bound Rof sterically hinders RNA accommodation at the extended PBS. (F) Ring closure. Left, fluorescence anisotropy of 5’-FAM-labeled rU_12_ in the presence of increasing amounts of ρ. Right, Rof inhibits ρ ring closure. Data of two independent experiments and fits to a single-exponential Hill function (see Materials and Methods) are shown.

**Figure 5.**
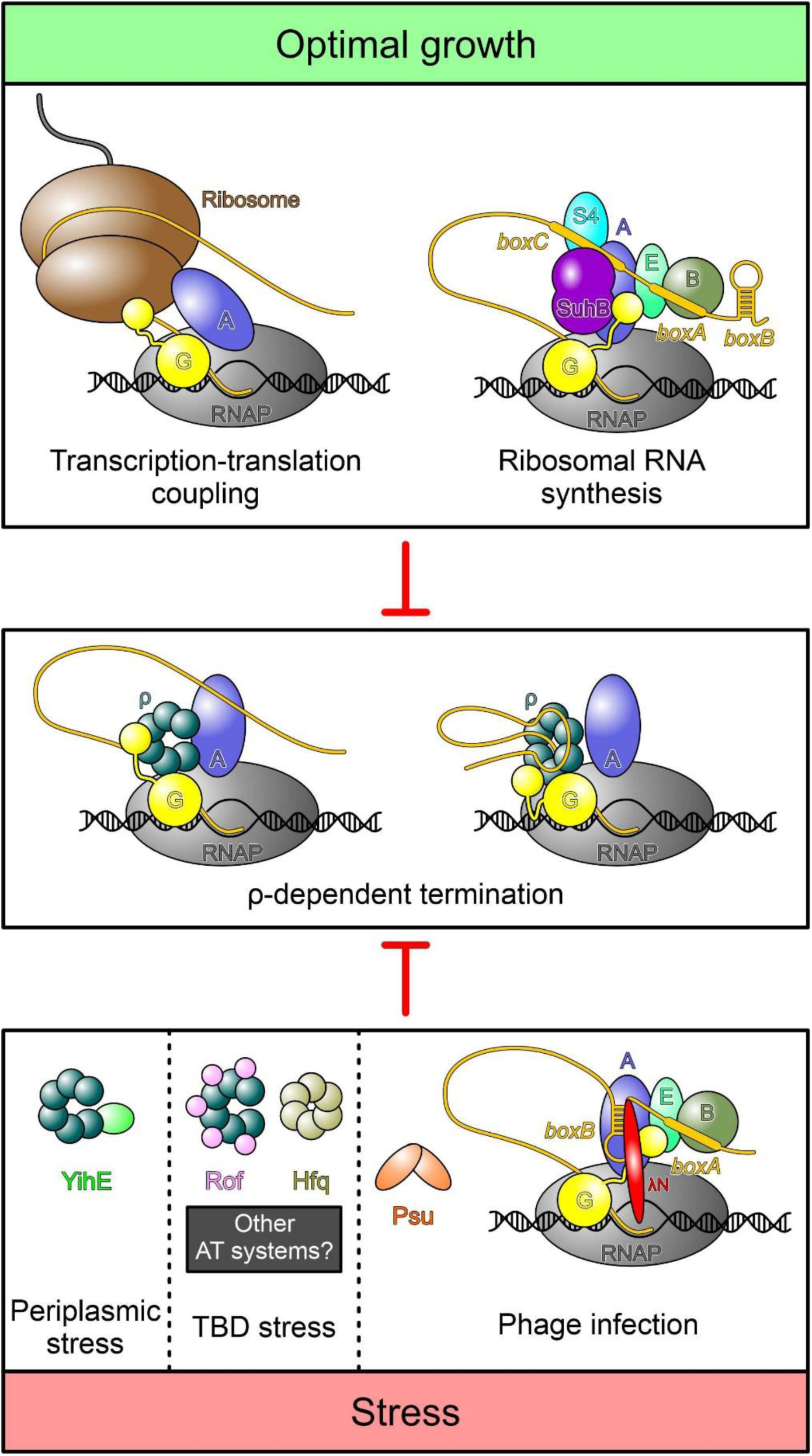
Top, under optimal growth conditions bulk mRNA synthesis is protected from premature termination by ρ through transcription-translation coupling (65, 66). A specific transcription anti-termination complex (*rrn*TAC) shields the ribosomal RNAs from ρ (10). Bottom, under stress conditions, when translation is inefficient, ρ activity must be regulated. Phages use different strategies to protect transcription of their own genomes. Similar to the *rrn*TAC, lambdoid phages rely on specific anti-termination complexes, such as λN-TAC, to shield the nascent transcript from ρ (11). Phage P4 uses the Psu protein that directly binds to ρ and thereby prevents the formation of termination complexes (36). To date, three cellular proteins are known to directly bind ρ and inhibit termination, but other stress-specific regulators likely exist. YihE is induced upon periplasmatic stress (43, 51), whereas stress conditions under which Hfq and Rof regulate ρ activity remain to be identified. YihE binds the ρ NTD, inhibiting ρ-RNA interactions (20); the mechanism of ρ regulation by Rof has been determined in this study, and molecular details of Hfq-mediated ρ inhibition remain to be solved (16, 19). By directly binding ρ, each of these anti-terminators may interfere with the formation of ρ-dependent termination complexes that assemble in an EC-dependent manner (center left) (31, 32) or RNA-dependent manner (center right) (47, 48).

Rof molecules are bound between protomers ρ_A_-ρ_B_, ρ_B_-ρ_C_, etc., but not preceding or following the terminal ρ subunits, ρ_A_ and ρ_F_ (Fig. 2*B*). In each case, Rof interacts primarily with one ρ NTD through its N-terminal α-helix (α1, residues 9-23), the loop connecting strands β3 and β4 (residues 46-50), and the very C-terminal region (Fig. 2*C*). In addition, loop^β3-β4^ of Rof contacts the N-terminal three-helix bundle of the adjacent ρ protomer (Fig. 2*C*), presumably explaining why stable Rof binding is only observed between neighboring ρ subunits but not at the terminal subunits of the open ring. The N-terminus and helix α1 of Rof are positioned in a cavity of the ρ NTD between ρ helix α4 (residues 83-89) and one flank of the ρ OB-fold (residues 95-120; Fig. 2*C*). Close contacts between the Rof N-terminus and ρ could explain why the addition of a His-tag blocks Rof activity *in vivo* (Fig. 1*B*). Loop^β3-β4^ of Rof is positioned atop helix α5 within the ρ NTD-CTD linker (the “connector”) and the loop between helix α2 and α3 of the adjacent ρ protomer (residues 22-30; Fig. 2*C*). The very C-terminus of Rof rests on the exposed surface of the ρ OB-fold next to the Rof N-terminal region (Fig. 2*C*).

Contacts between ρ and Rof are mediated predominantly by polar and charged interactions (Fig. 2*D-H*). The backbone carbonyl oxygen of Rof^I8^ (N-terminus) as well as the side chains of Rof^Y13^ (helix α1) and Rof^E50^ (β3-β4 loop) contact ρ^R88^ by hydrogen bond, cation-π and ionic interactions, respectively (Fig. 2*D*). Rof^C10^ and Rof^D14^ (helix α1) form hydrogen bonds with ρ^Q85^ and ρ^S82/S84^, respectively (Fig. 2*E*). Rof^E17^ (helix α1) forms a salt bridge to ρ^R87^ (Fig. 2*F*). Rof^N48^ (β3-β4 loop) forms a hydrogen bond with ρ^R128^ (connector), while Rof^K47^ interacts electrostatically with ρ^E24^ of the adjacent ρ subunit (Fig. 2*G*). The C-terminal carboxy group of Rof^S84^ is engaged by ρ^K100/K115^ (Fig. 2*H*). Most of the ρ-contacting residues of Rof are conserved (Fig. S6*A*). Likewise, Rof-contacting residues of ρ are highly conserved (Fig. S6*B*). As ρ is more widespread among bacteria than Rof, the conservation of the Rof-contacting ρ residues likely reflects their importance in forming the PBS.

### Replacement of ρ-contacting residues undermines Rof function

To validate our structural observations, we replaced Rof residues Y13, D14, or E17 with alanine and tested binding of the corresponding Rof variants to ρ^wt^ in analytical SEC (Fig. S7*A*). While the ρ interaction was only very mildly affected by an E17A exchange in Rof, ρ binding was completely abolished in Rof^Y13A^ and Rof^D14A^, as monitored by SEC. We also generated ρ variants, ρ^R88E^, ρ^F89S^, and ρ^K115E^, and tested them for Rof^wt^ binding (Fig. S7*A*). Interaction of ρ^R88E^ with Rof^wt^ was abrogated in SEC, while Rof^wt^ binding was only slightly reduced in ρ^F89S^ or ρ^K115E^. Thus, our structures delineate a precise network of charged and polar interactions that are crucial for ρ-Rof complex formation.

Rof variants that were unable to bind ρ were also unable to elicit anti-termination *in vitro*: ρ terminated synthesis of 90 % and 86 % of transcripts in the presence of Rof^Y13A^ and Rof^D14A^, respectively, as compared to 16 % in the presence of Rof^wt^ and 87 % in the absence of Rof (Fig. S7*B*). In agreement with only slightly reduced affinity of Rof^E17A^ for ρ, Rof^E17A^ was only slightly less effective in inhibiting ρ than Rof^wt^ (25 % of transcripts terminated in the presence of Rof^E17A^; Fig. S7*B*). These results indicate that the ρ-Rof contacts observed in our structures are relevant for Rof-dependent inhibition of ρ in the context of complete ECs.

Next, we evaluated the effects of Rof substitutions on toxicity *in vivo*. We measured the lag phase extension upon IPTG induction of plasmid-encoded Rof variants (Fig. S7*C*). While Rof^wt^ increased the lag 4.6-fold, substitutions of several conserved residues in the N-terminal helix of Rof strongly reduced this effect: plasmids guiding the production of Rof^Y13A^, Rof^D14A^, or N-terminally His-tagged Rof were indistinguishable from the empty vector, whereas Rof^C10S^ increased the lag 2.4-fold. As Rof^E17A^ did not show significant defects *in vitro* (Fig. S7*B*), we tested the effects of the Rof^E17K^ variant instead; the charge reversal led to a complete loss of Rof toxicity. By contrast, substitutions in the Rof β3-β4 loop (Fig. 2*G*) had smaller effects: R46A and K47A exchanges in Rof increased the lag 2.6- and 3.2-fold, respectively, and Rof^N48A^ was almost indistinguishable from Rof^wt^ (Fig. S7*C*). We conclude that Rof residues that engage in contacts to ρ in our structures are important for Rof-mediated inhibition of *E. coli* growth.

Substitutions of the surface-exposed Rof residues that contact ρ would not be expected to alter the Rof structure, and thus cellular production or stability. To ascertain that the observed *in vivo* defects are due to the loss of binding of the Rof variants to ρ, we used Western blotting to quantify Rof variants in cells. As specific anti-Rof antibodies are not yet available, we inserted a single HA tag into a flexible loop of Rof at residue 57, the only location that we found to tolerate tags without the loss of *in vivo* activity. We found that D14A, E17K, R46A and E47A variants of Rof were produced at levels comparable to, or higher than, Rof^wt^, whereas the Rof^Y13A^ variant was less abundant (Fig. S7*D*).

### Rof undergoes pronounced conformational changes upon ρ binding

A solution structure (PDB ID: 1SG5) (21) and an AlphaFold (34) model of isolated *E. coli* Rof showed marked differences and high flexibility in the N-terminal region compared to our ρ-bound Rof. In the isolated state, the N-terminal region lacks contacts to the globular portion of Rof and thus remains flexible, while it is immobilized and caps one face of the OB-fold in the ρ-bound state (Fig. S8). Concomitantly, helix α1 is oriented differently relative to the β-barrel in the isolated and ρ-bound states (Fig. S8). As a consequence, the N-terminal region and helix α1 are repositioned relative to the other main ρ-interacting regions of Rof, the β3-β4 loop and the C-terminus (Fig. S8). The above comparisons show that Rof undergoes pronounced conformational changes and folding/immobilization transitions upon ρ engagement, which are required to generate the productive ρ-binding surface, composed of N-terminal region, helix α1, β3-β4 loop and C-terminus. In contrast to Rof, no global conformational changes are observed in the ρ protomers upon Rof engagement (e.g. compared to an open-ring, ANPPNP-bound ρ structure; PDB ID: 1PVO) (2).

A model of an *E. coli* ρ-Rof complex has previously been proposed based on the solution structure of Rof and NMR-guided docking (21). While the general binding site of Rof on the ρ NTDs agrees with our cryoEM structures, the previous docking model differs significantly in detail, as the conformational changes upon binding were not captured. Presently, we cannot exclude that a minor population of isolated Rof adopts a conformation observed in the ρ-bound state, and that binding might, thus, occur *via* a conformational selection mechanism. However, presently available data favor a mechanism that involves binding *via* a pronounced induced fit in Rof.

Irrespectively, apart from steric hindrance (see above), an N-terminal tag on Rof might also interfere with the conformational changes and folding/immobilization upon binding required for stable complex formation with ρ, as also observed with other proteins (35).

### Rof blocks an extended PBS of ρ

Electrophoretic mobility shift assays indicated that Rof interferes with RNA binding to the ρ PBSes (21). To further evaluate Rof effects on RNA binding at ρ PBSes, we quantified the affinity of a 5’-FAM-labeled DNA oligomer, dC_15_, that binds exclusively to the ρ PBS (6) by fluorescence anisotropy (Fig. 3*A*). ρ alone bound dC_15_ tightly (*K_d_* = 1 µM; Fig. 3*B*), in agreement with previous results (36). ρ binding to dC_15_ was reduced stepwise in the presence of increasing concentrations of Rof^wt^, but not in the presence of Rof^Y13A^ or Rof^D14A^ (Fig. 3*B*), suggesting that bound Rof competitively or allosterically occludes the ρ PBSes or parts thereof.

A structure of ρ bound to a short oligonucleotide at the PBS (PDB ID: 1PVO) had delineated a core PBS that accommodates a dinucleotide (2). Comparison to the ρ-Rof complex structures showed that Rof binds next to the dinucleotide and does not directly interfere with RNA contacts at the core PBS (Fig. S9*A*). However, biochemical studies had suggested that more than a dinucleotide contributes to the affinity of RNA for the ρ NTD (37, 38). Furthermore, longer PBS ligands bind ρ more tightly and have greater impact on ring closure (3, 6). We thus sought to trace the complete RNA path along the ρ NTD. To this end, we solved the cryoEM/SPA structure of ρ in complex with a 99-nt long, natural *rut* RNA derived from the λ *tR’* terminator region (Fig. S9*B*). In the presence of ADP-BeF_3_ we obtained one reconstruction at a global resolution of 2.9 Å (Fig. S10 and S11), in which ρ adopted a closed ring conformation (Fig. 3*C*), in agreement with previous results that RNA and ATP jointly trigger ρ ring closure (3, 6).

The structure of ρ in the ρ-*rut* RNA complex is very similar to a closed-ring structure of ρ obtained in complex with an rU7 SBS ligand and ADP-BeF_3_ (PDB ID: 5JJI) (3). Thus, the ρ protomers A-E in the ring are arranged as a smoothly ascending spiral staircase, and protomer F represents a “transition” subunit to close the ring. Nine nucleotides (nts) of single-stranded RNA bound at the SBS in the center of the ring are clearly defined in the cryoEM density (Fig. S9*C*). Protomers ρ_A_-ρ_E_ contact the SBS RNA *via* their Q-loops (residues 278-290) and R-loops (residues 322-326), while ρ_F_ lacks direct contacts to the SBS RNA; most direct contacts are non-sequence specific, involving backbone functionalities of both, ρ and the SBS RNA (Fig. S9*D*).

The cryoEM density also revealed regions of the *rut* RNA bound at the PBSes of all ρ subunits (Fig. 3*C*). Density corresponding to PBS-bound RNA regions was not equally well defined at each subunit, most likely due to the different fractions of C-residues in the bound RNA regions. Furthermore, no density was observed for RNA regions connecting the PBS-bound elements or between one PBS-bound RNA region and the SBS-bound RNA (Fig. 3*C*).

The limited local resolution precluded an unequivocal sequence assignment for PBS-bound RNA regions. The *rut* RNA we used for structural analysis contains two C-rich regions, *rutA* and *rutB*, and an intervening stem-loop structure (*boxB*; Fig. S9*B*). We observed a globular patch of density at the ρ_C_-ρ_D_ interface (Fig. 3*C*) that might correspond to the *boxB* stem-loop structure, which would map the canonical *rut* elements of λ *tR’* at the PBSes of ρ_C_, ρ_D_, and ρ_E_. Guided by this landmark, we only explicitly modeled a 6-nt portion of *rutA* bound at the PBS of ρ_C_ (Fig. 3*C,D* and S9*E*). In the resulting model, the two 3’-most nts of the bound region are accommodated at the canonical core PBS, contacting ρ^F64^ and ρ^Y80^. The path of the preceding four nucleotides is lined by residues ρ^K102^, ρ^R105^, and ρ^K115^ on one side, and by residues ρ^D60^, ρ^F62^, ρ^P83^, ρ^S84^, ρ^R87^, and ρ^R88^ on the other. The 5’-terminal nt of the bound RNA region is embedded in a pocket formed by residues ρ^Q85^, ρ^R87^, ρ^R88^, ρ^F89^, and ρ^K115^ (Fig. S9*E*), with ρ^Q85/K115^ reaching towards the nucleobase (cytosine in our model) and ρ^R87/R88^ contacting the sugar-phosphate backbone. Consistent with our modeling, a cytidine at the 5’-end of an 8-nt long RNA oligomer increased the affinity of the RNA ligand to the ρ PBS 10-fold compared to a uridine, while purine nucleotides at this position completely abolished RNA binding (38).

To further test the importance of the additional RNA-binding regions of the ρ NTD for the interaction with a long *rut*-site RNA, we tested binding of the ρ^R88E^, ρ^F89S^, and ρ^K115E^ variants to the λ *tR’ rut* RNA by analytical SEC (Fig. S9*F*). While ρ^F89S^ did not show decreased binding to the RNA as compared to ρ^wt^, RNA binding was significantly reduced for ρ^K115E^ and completely abolished for ρ^R88E^. Taken together, our structural analysis revealed additional RNA-contacting regions in the ρ NTD that we refer to as the “extended PBS”, which augments the RNA affinity and specificity of the ρ core PBS.

Structural comparisons between our ρ-Rof and ρ-*rut* RNA structures showed that bound Rof would block RNA binding at the extended PBS (Fig. 3*E*). Consistent with the idea that Rof blocks ρ activity by outcompeting RNA at the ρ extended PBS, residues that contribute to the extended PBS are also important for Rof binding (Fig. 2*D-H* and Fig. 3*E*).

### RNA binding at the extended PBS supports ρ dynamics accompanying ring closure

To activate ATP hydrolysis and RNA translocation, ρ has to convert from an open to a closed-ring conformation (3, 4). Ring closure involves conformational changes in ρ that affect the protomer interfaces (3, 5). Upon ring closure, the NTDs of adjacent ρ subunits separate, breaking contacts between the NTD-CTD connector of one ρ protomer and the N-terminal three-helix bundle of the adjacent ρ protomer; simultaneously, the ρ CTDs approach each other more closely, encircling the RNA at the SBS and forming ATPase-active ATP-binding pockets (2, 5).

In open ρ structures, ρ^R128^ at the C-terminus of connector helix α5 contacts ρ^N25^ of a neighboring NTD, helping to position ρ^R28^ of the adjacent protomer in a binding pocket that involves helix α5 of the connector (Fig. S12*A*). These inter-subunit contacts are broken during ring closure; ρ^R128^ now hydrogen bonds to the backbone carbonyl of ρ^R88^ and/or is engaged by the side chain of ρ^E125^, and ρ^R28^ is exchanged for ρ^K130^ of the same protomer (Fig. S12*B*). Furthermore, in open and closed ρ structures, the NTD-CTD connector establishes different contacts to the NTD and CTD of its own protomer as well as to the neighboring ρ subunit; upon ring closure, interactions along the entire NTD-CTD connector shift in register by one residue (39, 40) (Fig. S12*C*). Thus, the NTD-CTD connector establishes dynamic interaction networks, differentially interconnecting the NTD and the CTD within a protomer as well as differentially contacting the neighboring protomer in the open and closed states.

Apart from ATP binding between neighboring CTDs and RNA binding at the SBS, ring closure is also promoted by RNA binding at the PBSes (25, 40). However, available structures of ρ in complex with PBS ligands failed to uncover the structural basis for this effect. RNA binding at the extended PBS, as observed in our ρ-*rut* RNA structure, could solve this riddle. Residues forming the extended PBS and the 5’-end of the PBS-bound RNA region are in direct proximity of connector helix α5 (Fig. S12*D*). We surmise that RNA bound at the extended PBS supports the closed-ring interaction network by structurally stabilizing the NTD, reinforcing the sequestration of ρ^R128^ and the repositioning of ρ^K130^ (Fig. S12*B,D*). It is also conceivable, albeit not resolved in our ρ-*rut* RNA structure, that the 5’-end of RNA regions bound at the extended PBS directly contacts the neighboring connector helix α5. PBS-bound RNA could thereby stabilize the connector in the closed-ring configuration and support an apparent rotational movement of the NTD, which facilitates displacement of the three-helix bundle from the neighboring NTD, thus supporting retraction of ρ^R28^ from its inter-subunit binding pocket (Fig. S12*A,B*).

### Rof stabilizes the open ρ conformation and undercuts connector-mediated communication between ρ domains

Upon binding, Rof positions its β3-β4 loop on top of helix α5, establishing direct contacts of Rof^N48^ to ρ^E125^ or ρ^R128^ and thus reinforcing the interaction of ρ^R128^ with ρ^N25^ of the neighboring NTD (Fig. S12*E*). Furthermore, Rof^K47^ in the β3-β4 loop engages in a salt bridge with ρ^E24^ of the neighboring ρ protomer (Fig. 2*G* and S12*E*). Thus, Rof bound between two neighboring NTDs of open ρ may stabilize their relative positions, and thus the open-ring conformation. To test whether Rof indeed counteracts ρ ring closure, we performed fluorescence anisotropy assays with the 5’-FAM-labeled SBS ligand, rU_12_, in the presence of unlabeled dC_15_ that blocks the PBSes (6), such that rU_12_ binding at the SBS reports on ρ ring closure (Fig. 3*A*). Indeed, while ρ alone readily underwent ring closure, Rof inhibited ring closure in a concentration-dependent manner (Fig. 3*F*). Surprisingly, although ρ adopts an open conformation upon Rof binding, the connector retains the NTD/CTD-interaction register of the closed conformation (Fig. S12*C*), presumably due to the direct Rof-helix α5 interactions restricting connector rearrangements. Concomitantly, in complex with Rof, the CTD Q-loops adopt a conformation observed in closed ρ, but not in open ρ structures in the absence of Rof (41) (Fig. S12*F*); in the closed ring as well as in the open ring in complex with Rof, ρ^K283^ is inserted into a pocket formed by the Q-loop of the adjacent protomer, while ρ^K283^ points to the center of the ring in open ρ in the absence of Rof (Fig. S12*F*).

Together, these observations show that Rof effectively stabilizes ρ in an open conformation by cross-strutting neighboring NTDs and preventing their separation, yet at the same time prevents a register shift and change in interactions along the NTD-CTD connector, maintaining the CTDs in a conformation resembling the closed state. Thus, Rof conformationally insulates ρ NTDs and CTDs by undercutting concerted structural transitions normally associated with ρ ring dynamics and induces a hybrid conformation, with the NTDs in the open state and the connector and CTDs in the closed-state conformation.

### Rof interferes with ρ binding to transcription elongation complexes

Recent structural studies had shown that ρ can engage ECs modified by general transcription factors NusA and NusG before binding the transcript, with the ρ NTDs establishing direct contacts to NusA, NusG and RNAP subunits α, β and β’ (31, 32). In the ρ-bound NusA/NusG-ECs, ρ remains dynamic relative to the surface of RNAP, allowing ρ to position one of its PBS next to the RNA exit channel to eventually capture the nascent transcript. Subsequent conformational changes in RNAP can trigger transcription inactivation.

Superposition of the ρ-Rof and ρ-EC structures revealed that ρ surfaces in contact with ECs encompass the extended PBSes, and that Rof binding to ρ is incompatible with ρ binding to an EC (Fig. S13*A*). Consistently, when in complex with Rof, ρ showed impaired binding to a NusA/NusG-EC in analytical SEC (Fig. S13*B*). We observed partial dissociation of the ρ-Rof complex in favor of ρ binding to the EC, consistent with a competition between Rof and EC for ρ. Similarly, ρ dissociates partially from a pre-formed ρ/NusA/NusG-EC upon addition of Rof, indicating that Rof can act on ρ-bound transcription complexes within the cell. Thus, apart from preventing productive engagement of a transcript, Rof hinders EC engagement by ρ.

### Rof proteins from *Vibrionaceae* may resort to a modified ρ-inhibitory mechanism

Unlike ρ, which is ubiquitous in bacteria, Rof is restricted to Pseudomonadota (synonym Proteobacteria; Fig. 1*D*); 79 % of *Enterobacteriaceae* and 63 % of *Vibrionaceae* genomes are estimated to encode Rof (Dataset S1*A*). In *E. coli*, *rof* is encoded in a five-gene locus bracketed by REP elements (Fig. 1*E*). Two genes in this cluster, *arfB* and *nlpE*, play vital roles in *E. coli* stress response. ArfB, a peptidyl-tRNA hydrolase, rescues stalled ribosomes when trans-translation control fails (42). NlpE is an outer membrane lipoprotein that activates the Cpx pathway in response to envelope stress (43). Rof is expressed from a divergent promoter in tandem with a small conserved protein of unknown function, YaeP (44).

Presence of *rof* in diverse Pseudomonadota (Fig. 1*D*) suggests a common mechanism of ρ inhibition. However, recombinant *V. cholerae* Rof (*vc*Rof) has been observed to form higher-order oligomers *in vitro*, of which the dimer state could be stabilized by treatment with iodoacetamide/TCEP; furthermore, *vc*Rof seems to disassemble ρ hexamers *in vitro* (22). ρ-bound *E. coli* Rof (*ec*Rof) and a *vc*Rof monomer (PDB ID: 6JIE) (22) exhibit similar structures that superimpose with a root-mean-square deviation (rmsd) of 1.05 Å for 58 pairs of Cα atoms (2.65 Å for all 72 pairs of Cα atoms). Similar to isolated *ec*Rof, isolated *vc*Rof differs from ρ-bound *ec*Rof in the orientation of the N-terminal region, which is diverted from the OB-fold; furthermore, helix α1 in isolated *vc*Rof is shortened by one turn compared to *ec*Rof (Fig. 4*A,B*). Given that crucial ρ-contacting residues in the N-terminal regions are highly conserved between *ec*Rof and *vc*Rof (Fig. 4*C*), *vc*Rof most likely binds its cognate ρ in a similar manner and by undergoing similar conformational changes as revealed here for *ec*Rof. However, more pronounced sequence divergence in other regions of Rof proteins from *Enterobacteriaceae* and *Vibrionaceae* (Fig. 4*C*), including in the ρ-contacting β3-β4 loops (Fig. 2*G*), may reflect some differences in the specific contacts to ρ.

**Figure 4.**
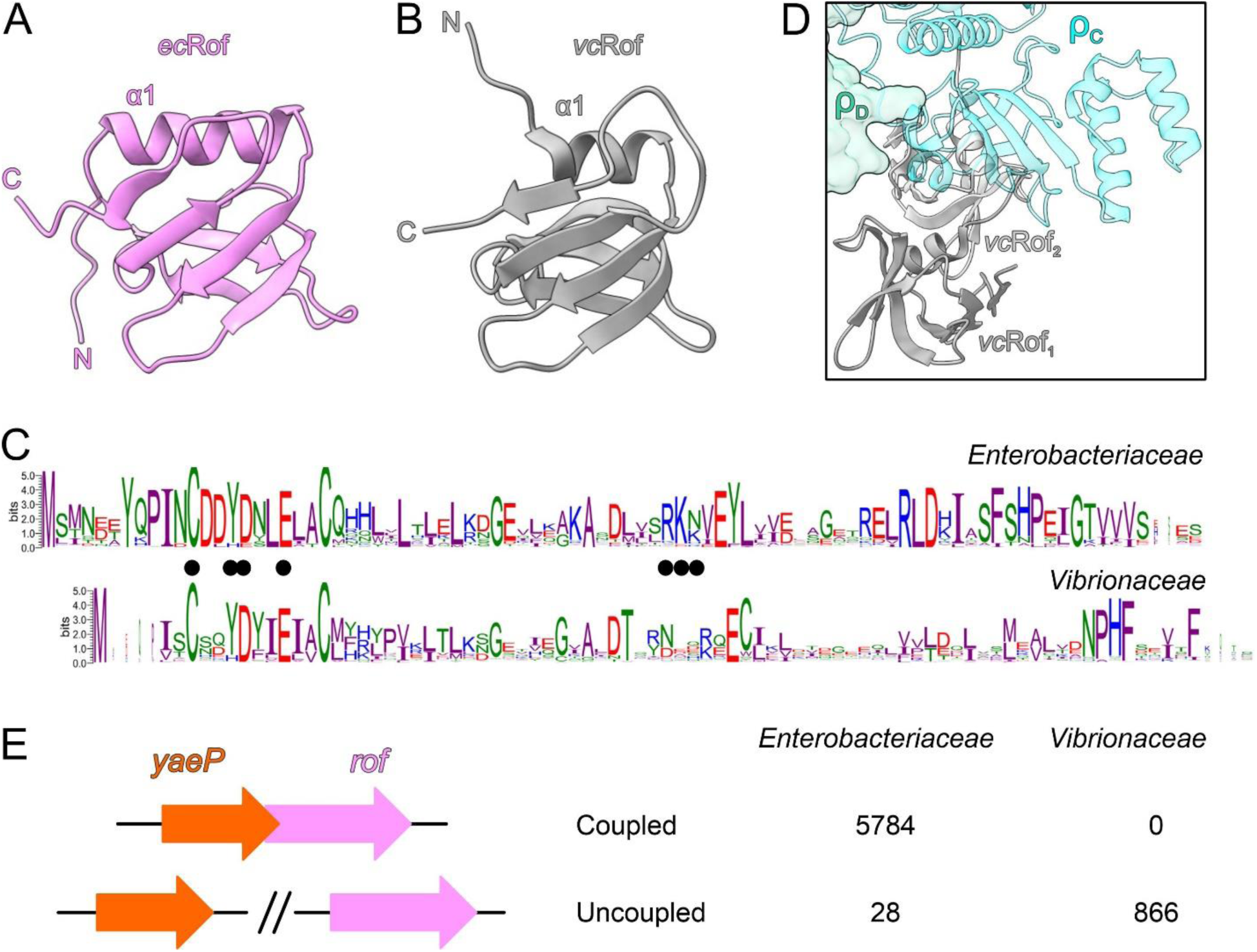
(*A*, *B*) The structure of one *E. coli* (*ec*) Rof protein as observed in the ρ-Rof complexes (*A*) compared to one monomer from the crystal structure of a *V. cholerae* (*vc*) Rof dimer (*B*) (PDB ID: 6JIE). *Ec*Rof bound to ρ and *vc*Rof mainly differ in the orientation of the N-termini and the lengths of helices α_1_. N-termini, N; C-termini, C. (A) (C) Conservation of Rof sequences in *Enterobacteriaceae* (top) and *Vibrionaceae* (bottom). In *Enterobacteriacea*, two alternative start sites for Rof are possible, generating 84- and 86-residue Rof. The sequence logos were generated by WebLogo (version 3.7.8). The interface residues revealed by the *E. coli* ρ-Rof complex structures are indicated by black dots. (B) (D) Superposition of the *vc*Rof dimer on *ec*Rof bound to ρ. While monomeric *vc*Rof would align without steric conflict, there might be steric hindrance in the interaction between a dimer of *vc*Rof and ρ. (C) (E) The *rof* and *yaeP* genes are coupled in *Enterobacteriaceae*, but not in *Vibrionaceae*; see Dataset S1*B*,*C* for more details.

A heterotypic configuration of the *vc*Rof dimer has been proposed based on a crystal structure (PDB ID: 6JIE) (22). While putative ρ-interacting surfaces, in particular the N-terminal region, of one *vc*Rof subunit are involved in the proposed dimerization, the putative ρ-interacting elements of the other *vc*Rof subunit are unobstructed; thus, while the proposed *vc*Rof dimer could engage ρ *via* one of the *vc*Rof subunits as observed for our *E. coli* ρ-Rof complexes without steric hindrance, conflicts would ensue upon dimeric *vc*Rof binding to ρ *via* the subunit with partially buried ρ-interacting elements (Fig. 4*D*). While such engagement may in principle lead to conformational changes in ρ and disassembly of the hexamer, the molecular mechanisms that would favor this disruptive engagement over binding of dimeric *vc*Rof *via* the unobstructed subunit remain unclear.

Apart from differences in the interaction details and the different oligomeric states, other factors may differentially influence the mode of action of Rof proteins in *Enterobacteriaceae* and *Vibrionaceae*. In agreement with differential regulation by the cellular environments, we observed that the *rof* genomic contexts in *Enterobacteriaceae* and *Vibrionaceae* are strikingly different. While in *Enterobacteriaceae,* the *rof* gene always overlaps with an upstream *yaeP* gene, the *yaeP* and *rof* genes are located on different chromosomes in *V. cholerae* (Fig. 4*E* and Dataset S1*B*,*C)*. Furthermore, *rof* does not have conserved gene neighbors in *Vibrionaceae* (Fig. S14 and Dataset S1*D*). Collectively, our analyses reveal that monomeric *ec*Rof and dimeric *vc*Rof could engage ρ in an analogous fashion and thus inhibit ρ function *via* similar molecular principles. The molecular mechanisms of putative additional levels of ρ inhibition *via* subunit dissociation by *vc*Rof as well as possible differences in Rof regulation remain to be elucidated.

## Discussion

ρ is a global gene regulator and essential in most bacterial species. It defines gene boundaries and silences production of useless and harmful RNA. *In vitro*, ρ-dependent termination can proceed *via* two pathways (45, 46). In the RNA-dependent pathway, ρ binds C-rich, unstructured and unoccupied RNA regions *via* its PBSes; subsequent binding of a neighboring RNA region at the SBS and ring closure allow ρ to translocate towards elongating RNAP, eventually contact the transcriptional machinery around the RNA exit channel and terminate transcription (47, 48). In the EC-dependent pathway, ρ associates with and accompanies ECs without immediately terminating transcription (49); upon EC pausing, NusA and NusG can cooperate with ρ in inducing conformational changes in RNAP that stall transcription before ρ engages the transcript (31, 32). Under stress conditions when dedicated anti-termination machineries and translating ribosomes are scarce, ρ action may need to be tuned down. To date, three cellular proteins that inhibit ρ function, Hfq, YihE and Rof, have been described (19–21, 50). While a recent study revealed molecular details for YihE (20), the mechanisms of ρ inhibition by the Sm-like proteins Rof and Hfq remain incompletely understood. Here we present structural, functional, and phylogenetic analyses of ρ regulation by Rof. Extending and detailing an earlier docking model (21), our results show that Rof undergoes pronounced conformational changes and folding/immobilization upon binding to the NTDs of an open ρ ring, thereby implementing multiple strategies to counteract both ρ-dependent termination pathways.

First, Rof prevents ρ from binding to RNA at the PBS by blocking an extended PBS. An extended PBS has long been envisioned to contribute to both the RNA affinity and specificity of ρ, even though structures of open ρ revealed only a dinucleotide bound at the canonical PBS (PDB ID: 1PVO) (2), and structures of closed ρ lacked RNA at the PBS (PDB ID: 3ICE) (5). Recent structures of ρ in complex with transcribing RNAP showed RNA bound at each of the PBSes and at the SBS of a closed ρ ring (47, 48) in a manner similar to our ρ-*rut* RNA structure; however, limited resolution in the ρ portions of the structures precluded unequivocal assignment of the extended PBS. Our ρ-*rut* RNA structure clearly delineated the extended PBS (Fig. 3*D*). Comparison to our ρ-Rof structures revealed that ρ residues contacting Rof are also crucial for binding RNA at the extended PBS (Fig. 3*E* and Fig. S9) and are highly conserved (Fig. S6*B*).

Second, Rof prevents ρ ring closure and thus RNA binding at the SBS (Fig. 3*F*). Given that optimal SBS ligands (*e.g.*, rU_12_) promote ring closure even in the absence of a PBS ligand (6), PBS blocking alone cannot fully accounts for Rof-mediated inhibition of ρ ring closure. Furthermore, Rof itself is a PBS ligand – why does Rof binding not induce ring closure? Our results show that Rof bridges the NTDs of two adjacent ρ protomers, thus preventing their separation that is required for ring closure (Fig. 2*C,G*).

Third, Rof conformationally insulates the ρ NTDs and CTDs, undercutting conformational dynamics involved in RNA binding at the SBS and ring closure. Functional studies had suggested that residues forming the core and extended PBSes are involved in an allosteric network that mediates communication across ρ domains *via* the NTD-CTD connector. For example, a Y80C exchange in the NTD alters the CTD conformation (39), and ρ^R88^ has been suggested to affect steps other than RNA binding at the PBS (37). Our ρ-*rut* RNA structure suggests that such communication involves stabilization of the bound regions, which strengthens contacts of the NTD-CTD connector characteristic of the closed state. In addition, the 5’-end of PBS-bound RNA could directly contact the connector helix α5 and stabilize the closed-state configuration. Rof, in contrast, induces a chimeric structure in ρ. While in the presence of Rof NTDs adopt an open-ring arrangement, interaction patterns and conformations of the NTD-CTD linker and the CTDs resemble the closed state, presumably due to Rof directly contacting helix α5 of the connector, thus stabilizing the closed-state register. As a consequence, in the presence of Rof, the CTDs remain in a conformation resembling the closed-state; in particular, the Q-loops adopt a conformation very similar to that observed in closed-ring structures of ρ, with ρ^K283^ embedded in a pocket formed by the Q-loop of the neighboring protomer. In open ρ complexes in the absence of Rof, ρ^K283^ side chains instead point to the ring interior (PDB ID: 6WA8) (41). We speculate that ρ^K283^ in open ρ might be important for the initial RNA capture at the SBS and its proper alignment during ring closure. Thus, by retaining CTDs in a state resembling the closed ring, and thereby sequestering ρ^K283^, Rof might also counteract initial RNA engagement at the SBS.

Fourth, as the extended PBSes and Rof-contacting surfaces on ρ also represent main contact points of ρ on NusA/NusG-modified ECs (31, 32), Rof prevents ρ from interacting with ECs (Fig. S13). Rof interferes with ρ binding to the EC and is able to partially dissociate ρ from pre-formed ρ-ECs (Fig. S13*B*).

Rof-mediated inhibition of ρ complements other strategies of ρ-regulation in bacteria (Fig. 5). Maintaining a balance between inactive ρ sequestered by Rof and unobstructed ρ that could engage nascent transcripts or ECs, would allow cells to tune ρ-dependent transcription termination in response to cellular cues, e.g., during slow growth or under stress. At least two key questions concerning the cellular role of Rof presently remain unanswered: When does the cell need Rof and how is Rof expression controlled? Rof is present in many Pseudomonadota, implying that its ability to dampen ρ-dependent termination is important under some physiological conditions, but the *rof* deletion in *E. coli* has no known growth phenotypes. As ρ is essential, a global anti-terminator should be produced in substantial amounts only when needed. Indeed, a quantitative proteomic analysis of *E. coli* (17) demonstrated that both Rof and YihE are present at low (and similar) levels across many conditions. Consistent with published data (23), Rof levels modestly increased during the stationary phase, and a similar trend was observed for YihE (17). The signal that activates YihE is known: YihE is regulated by Cpx (51), a two-component system that responds to envelope stress (43). Collectively, it stands to reason that the expression of *yaeP-rof* is induced by particular cellular signals; e.g., a modest increase in the *yaeP-rof* transcript level was observed during recovery from exposure to mercury (52). Identification of other triggers awaits a systematic analysis. Bacteria experience a wide variety of adverse conditions and inhibiting ρ could be essential for survival in some, but not other, situations. We hypothesize that Rof and YihE belong to a large set of specialized anti-termination factors, which could comprise proteins and sRNAs, that turn off ρ under diverse stress conditions to enable survival and facilitate recovery.

## Materials and Methods

### Recombinant protein and RNA production and purification

Table S1 lists plasmids used in this study. *E*. *coli* ρ protein and variants thereof were produced and purified as described before (11). DNA encoding Rof was codon optimized for *E. coli* expression and purchased from Invitrogen (Thermo Fisher Scientific). The *rof* gene was cloned into pETM-11 and expressed in *E*. *coli* RIL (BL21) cells (Novagen). Cells were lysed by sonification in buffer A (100 mM KCl, 5 mM MgCl_2_, 20 mM Na-HEPES, pH 7.5, 1 mM DTT, 10 % [v/v] glycerol, 1 mM DTT) and the lysate was cleared by centrifugation at 21,500 rpm for 1 hour at 4°C. The cleared lysate was loaded on a 5 ml Ni^2+^-NTA column (Cytiva) equilibrated in buffer A. Rof was eluted in elution buffer (buffer A supplemented with 500 mM imidazole). The eluate was treated with TEV protease and dialysed overnight against buffer A with 5 % (v/v) glycerol. After recycling on the Ni^2+^-NTA column, the flow-through was collected, concentrated, and loaded to a Superdex 75 column (Cytiva) equilibrated in SEC buffer (100 mM KCl, 5 mM MgCl_2_, 20 mM Na-HEPES, pH 7.5, 1 mM DTT). Peak fractions were pooled and concentrated to 4.4 mM. Rof was flash frozen in liquid nitrogen and stored at -80°C until further use. *rho* and *rof* mutations were introduced by site directed mutagenesis using the QuikChange protocol (Stratagene) and protein variants were produced and purified as for the wt versions. *In vitro* transcribed *rut* RNA was produced and purified as described before (31).

### SEC-MALS

SEC-MALS was performed in reaction buffer (20 mM Tris-HCl, pH 8.0, 120 mM KOAc, 5 mM Mg(OAc)_2_, 2 mM DTT). 1 mg/ml of ρ were mixed with 2 mg/ml of Rof and incubated at 32 °C for 10 min. 60 µl of the mixture were loaded on a Superose 200 increase 10/300 GL column (Cytiva) and chromatographed on HPLC system (Agilent) coupled to a miniDAWN TREOS multi-angle light scattering and a RefractoMax 520 detector system (Wyatt Technologies). Prior to measurements, a system calibration was performed using BSA (Sigma-Aldrich). ASTRA 6.1 software was used for data analysis and molecular mass determination.

### Sample preparation for cryoEM

For ρ-Rof complex formation, 41.8 µM ρ hexamer were mixed with a 10-fold molar excess of Rof in the absence or presence of 2.3 mM ADP-BeF_3_ in reaction buffer and incubated for 15 min at 32°C. The mixture was applied to a Superdex 200 Increase 3.2/300 column (Cytiva) and fractions of the complex were pooled and concentrated. 3.8 µl of the purified complex (5.3 mg/ml) were applied to glow-discharged Quantifoil R1.2/1.3 holey carbon grids and plunged into liquid ethane using a Vitrobot Mark IV (Thermo Fisher) set at 10 °C and 100 % humidity.

For ρ-*rut* RNA complex formation, 41.8 µM ρ hexamer were mixed with an equimolar amount of *rut* RNA and 2.3 mM ADP-BeF_3_ in reaction buffer and incubated for 15 min at 32 °C. The mixture was applied to a Superose 6 Increase 3.2/300 column (Cytiva) and fractions of the complex were pooled and concentrated. Purified complex (3.8 mg/ml) was vitrified as above.

### CryoEM data acquisition and processing

CryoEM data were acquired on a FEI Titan Krios G3i TEM operated at 300 kV equipped with a Falcon 3EC direct electron detector. Movies for ρ-Rof samples were taken for 40.57 s accumulating a total electron dose of ∼ 40 e^-^/Å^2^ in counting mode distributed over 33 fractions at a nominal magnification of 96,000 x, yielding a calibrated pixel size of 0.832 Å/px. Data for the ρ- *rut* RNA sample were acquired at a higher nominal magnification of 120,000 x, corresponding to a calibrated pixel size of 0.657 Å/px. A total electron dose of 40 e^-^/Å^2^ was accumulated over an exposure time of 30.58 s.

All image analysis steps were done with cryoSPARC (version 4.1.1) (53). Movie alignment was done with patch motion correction, CTF estimation was conducted with Patch CTF. Class averages of manually selected particle images were used to generate an initial template for reference-based particle picking from 4,216 micrographs for the ρ-Rof sample. 1,782,459 particle images were extracted with a box size of 384 px and Fourier-cropped to 96 px for initial analysis. Reference-free 2D classification was used to select 1,101,769 particle images for further analysis. *Ab initio* reconstruction was conducted to generate an initial 3D reference for heterogeneous 3D refinement. 600,888 particle images were further classified by 3D variability analysis into ρ_6_-Rof_5_ and ρ_5_-Rof_4_ complexes. Particle images were re-extracted with a box size of 384 px after local motion correction and subjected to non-uniform refinement giving reconstructions at ∼ 2.9 Å resolution each that could be slightly improved by CTF refinement. Another iteration of heterogeneous 3D refinement was applied to determine the final set of 110,264 particle images for the ρ_6_-Rof_5_ complex and 293,352 particle images for the ρ_5_-Rof_4_ complex. Non-uniform (NU) refinement yielded reconstructions at a global resolution of 2.92 Å and 2.74 Å, respectively. Data analysis for the ρ-Rof samples was conducted similarly.

### Model building, refinement, and analysis

Coordinates of ρ (PDB ID: 6WA8 [open ring]; PDB ID: 1PVO [closed ring]) and Rof (PDB ID: 1SG5) were docked into the cryoEM maps using Coot (version 0.8.9) (54). ρ subunits and Rof were manually adjusted to fit the cryoEM density. The *rut* RNA regions were built *de novo* except for the dinucleotide at the canonical PBS (taken from PDB ID: 1PVO) (2). Manual model building alternated with real space refinement in PHENIX (version 1.20.1) (55). Data collection and refinement statistics are provided in Table S2. Structure figures were prepared with ChimeraX (version 1.6.1) (56).

### Analytical size-exclusion chromatography

Proteins and/or nucleic acids were mixed in reaction buffer and incubated for at 32 °C for 10 min. 50 µl of the samples were loaded on a Superdex S200 P.C 3.2 column (ρ-Rof) or Superose 6 Increase 3.2/300 column (ρ-*rut* RNA) and chromatographed at a flow rate of 50 µl/min at 4 °C. To test for ρ-Rof interaction, 5 µM ρ and 90 µM Rof/variants were used. To test for ρ-*rut* RNA interaction, 4 µM ρ/variants and 5 µM *rut* RNA were used.

For testing ρ binding to ECs in the absence of presence of Rof, an EC composed of RNAP, NusA, NusG, nucleic acid scaffold without or with ρ was formed as described before (31). 2.4 µM EC lacking ρ were incubated with a 3-fold molar excess of ρ-Rof complex at 32 °C for 15 min. Alternatively, 1.1 µM ρ-EC were incubated with a 10-fold molar excess of Rof at 32 °C for 15 min. Fractions were analyzed by SDS-PAGE and urea PAGE to reveal protein and nucleic acid contents, respectively.

### Nucleic acid binding assays

Nucleic acid binding to ρ PBS or SBS was tested by fluorescence depolarization-based assays (6). For PBS binding, 5 µM 5’-FAM-labeled dC_15_ oligo (Eurofins) were mixed with increasing amounts of ρ (0 to 17 µM final hexamer concentration) in 20 mM Na-HEPES, pH 7.5, 150 mM KCl, 5 % (v/v) glycerol, 5 mM MgCl_2_, 0.5 mM TCEP. For SBS binding, ρ PBSes were first saturated with 10 µM non-labeled dC_15_. ρ (0 to 16 µM final hexamer concentration) was then mixed with 2 mM ADP-BeF_3_ and 2 µM 5’-FAM-labeled rU_12_ oligo (Eurofins). To test the effect of Rof and Rof variants on nucleic acid binding to ρ PBS or SBS, 1 or 5 µM of ρ hexamer were mixed with increasing amounts of Rof (0 to 100 µM) prior to dC_15_ addition. The fluorescence anisotropy was recorded in OptiPlateTM 384-well plates (PerkinElmer) using a Spark Multimode Microplate reader (Tecan; excitation wavelength, 485 nm; detected emission wavelength, 530 nm). Two technical replicates were averaged for each sample and the data were analyzed with GraphPad Prism software (version 9.0.2). To quantify ρ PBS or SBS binding, data were fitted to a single exponential Hill function; *Y* = *Y_max_*[protein]^h^ / (*K*_d_ ^h^ + [protein]^h^); *Y_max_*, fitted maximum of nucleic acid bound; *K_d_*, dissociation constant; h, Hill coefficient.

### Plating efficiency assays

Individual colonies of each strain, in 3 biological replicates, transformed with plasmids carrying inserts of interest under the control of P*_trc_* promoter were grown overnight in LB supplemented with carbenicillin at 32 °C. Serial dilutions of overnight cultures were spotted onto LB plates containing carbenicillin with and without IPTG.

### Growth assays

Individual colonies of each strain, in 4 biological replicates, were inoculated in MOPS EZ rich defined media (Teknova, #M2105) supplemented with carbenicillin (100 mg/l) and grown overnight at 32 °C. The overnight culture was diluted (1.5 μl into 98.5 μl) in fresh media supplemented with 1 mM IPTG, loaded into a 96-well plate, and grown in a BioTek EPOCH2 microplate reader at 32 °C. The OD_600_ in each well was measured at 15-min intervals for 24 hours with kinetic shaking and incubation. Results were analyzed in Microsoft Excel.

### Western blotting

Overnight cultures of cells transformed with plasmids encoding Rof variants were grown in LB + carbenicillin (100 mg/l) at 32 °C. After 1:100 dilution into fresh media, the cultures were grown to an early exponential phase at 32 °C, induced with 1 mM IPTG for 60 minutes, pelleted, and frozen. Cell pellets were resuspended in PBS, sonicated, and centrifuged. Protein concentration in cleared lysates was measured with Bradford reagent and normalized before resolving by SDS-PAGE. Proteins were transferred to a nitrocellulose membrane (Bio-Rad Trans-Blot Transfer Medium, Cat. 162-0112). Membrane was incubated with primary anti-HA antibody from mice (Sigma-Aldrich, Cat. H9658) and with secondary anti-mouse horseradish peroxidase linked antibodies (Amersham, Cat. NA931V) before imaging using ImageLab 3.0 with Bio-Rad ChemiDoc XRS+ system using Bio-Rad Clarity Max Western ECL Substrate (Cat. 1705062S).

### *In vitro* termination assays

RNAP and ρ for *in vitro* assays were purified as described in (31). RNAP holoenzymes were assembled by mixing the core RNAP with a three-fold molar excess of σ^70^ transcription initiation factor, followed by incubation at 30 °C for 20 min. DNA templates were generated by PCR amplification and purified by PCR cleanup kit (Qiagen). Halted A26 ECs were formed at 37 °C for 12 min by mixing 50 nM RNAP holoenzyme with 25 nM DNA template, 100 μM ApU, 10 μM ATP and UTP, 2 μM GTP, and 5 Ci/mmol [α^32^P] GTP in ρ termination buffer (40 mM Tris-Cl, 50 mM KCl, 5 mM MgCl_2_, 0.1 mM DTT, 3 % (v/v) glycerol, pH 7.9). After addition of ρ (25 nM) and Rof variants (5 μM), the reactions were incubated for 3 min at 37 °C. Transcription was restarted by addition of a pre-warmed (at 37 °C) mixture containing 150 μM NTPs and 50 μg/ml rifapentin and incubated at 37 °C for 7 min. Reactions were quenched by addition of an equal volume of stop buffer (45 mM Tris-borate, 10 M urea, 20 mM EDTA, 0.2 % xylene cyanol, 0.2 % bromophenol blue, pH 8.3) and separated by denaturing 5 % PAGE (7 M urea, 0.5X TBE). Gels were dried and products were visualized using a Typhoon FLA 9000 PhosphorImaging system (GE Healthcare). Readthrough and termination RNA products were quantitated with ImageQuant and Microsoft Excel software.

### Rof and ρ distribution in Pseudomonadota

Rof and ρ distribution (Dataset S1*A*) were estimated using Rof Pfam (PF07073) and ρ TIGR (TIGR00767) models in Annotree (57) with an e-value 10^-5^. The phylogenetic tree was downloaded from Annotree. The result was visualized in R (version 4.2.2) with ggtree package (58).

### Sequence conservation and genomic context analysis

To investigate the conservation of Rof and ρ in Pseudomonadota, the GTDB (59) bacterial taxonomy list (r_202) was downloaded. Only complete genomes were used to build the database. Three representatives from each Genus of Pseudomonadota were selected randomly. Rof was identified with the Pfam model. To identify ρ, Pfam model Rho_RNA_bind (PF07497) was searched against the representative database using hmmsearch (version 3.3) (60). The identified ρ-like proteins were further confirmed by searching NCBI HMM model (NF006886.1) against them using hmmsearch (60) (Dataset S1*E*,*F*).

To further investigate the conservation of Rof in *Enterobacteriaceae* and *Vibrionaceae*, reference genomes of *Enterobacteriaceae*, *Enterobacteriaceae*_A, and *Vibrionaceae* were selected from the GTDB bacterial taxonomy list. One representative from each strain was used to build the database. Again, Rof was identified by its Pfam model (Dataset S1*G*,*H*).

Before performing multiple sequence alignment, sequence duplicates were removed using cd-hit (version 4.8.1) (61). Sequences were aligned using MUSCLE (version 5.1) with default settings (62). Sequence logos were generated using WebLogo (version 3.7.8) (63).

*Enterobacteriaceae* and *Vibrionaceae* genomes containing *rof* and *yaeP* were analyzed to determine if their ORFs overlap; Dataset S1*B*,*C* is thus a subset of Dataset S1*G*,*H*. The overlaps were determined based on chromosomal location, strand, and gene distance. We note that instances of non-overlapping genes could be an artifact of low quantality genome assemblies.

### Data availability

CryoEM reconstructions have been deposited in the Electron Microscopy Data Bank (https://www.ebi.ac.uk/pdbe/emdb) under accession codes EMD-17875 (https://www.ebi.ac.uk/pdbe/entry/emdb/EMD-17875; ρ_6_-Rof_5_), EMD-17877(https://www.ebi.ac.uk/pdbe/entry/emdb/EMD-17877; ρ_5_-Rof_4_) EMD-17874 (https://www.ebi.ac.uk/pdbe/entry/emdb/EMD-17874; ρ_6_-ADP-Rof_5_), EMD-17876 (https://www.ebi.ac.uk/pdbe/entry/emdb/EMD-17876; ρ_5_-ADP-Rof_4_) And EMD-17870 (https://www.ebi.ac.uk/pdbe/entry/emdb/EMD-17870; ρ-*rut* RNA). Structure coordinates have been deposited in the RCSB Protein Data Bank (https://www.rcsb.org) with accession codes 8PTN (https://www.rcsb.org/structure/8PTN; ρ_6_-Rof_5_), 8PTP (https://www.rcsb.org/structure/8PTP; ρ_5_-Rof_4_), 8PTM (https://www.rcsb.org/structure/XXXX; ρ_6_-ADP-Rof_5_), 8PTO (https://www.rcsb.org/structure/8PTO; ρ_5_-ADP-Rof_4_) and 8PTG (https://www.rcsb.org/structure/8PTO; ρ-*rut* RNA). All other data are contained in the manuscript or the Supplementary Information. Source data are provided in this paper.

## Acknowledgments

We are very grateful to P. Lydia Freddolino and Natacha Ruiz for many stimulating discussions. This work was supported by the National Institutes of Health (GM067153 to I.A), the Deutsche Forschungsgemeinschaft (INST 130/1064-1 FUGG to Freie Universität Berlin; GRK 2473 "Bioactive Peptides", project number 392923329, to M.C.W.; WA 1126/11-1, project number 433623608, to M.C.W.) and the Berlin University Alliance (501_BIS-CryoFac to M.C.W.). The Sanger Sequencing at the OSU Comprehensive Cancer Center is supported in part by NCI P30 CA0168058.

## Author Contributions

N.S., B.W., T.L.S., D.G. and I.A. performed preparative and analytical biochemical experiments. M.F. and I.A. conducted *in vivo* experiments. B.W. carried out bioinformatic analyses. N.S. and T.H. prepared samples for cryoEM. T.H. collected cryoEM data and performed SPA. N.S. built and refined atomic models. All authors contributed to the analysis of data and the interpretation of the results. N.S. and I.A. wrote the manuscript with contributions from all authors. I.A. and M.C.W. coordinated the project.

## Competing Interest Statement

The authors declare no competing interests.

## Supplementary Information

**Figure S1.**
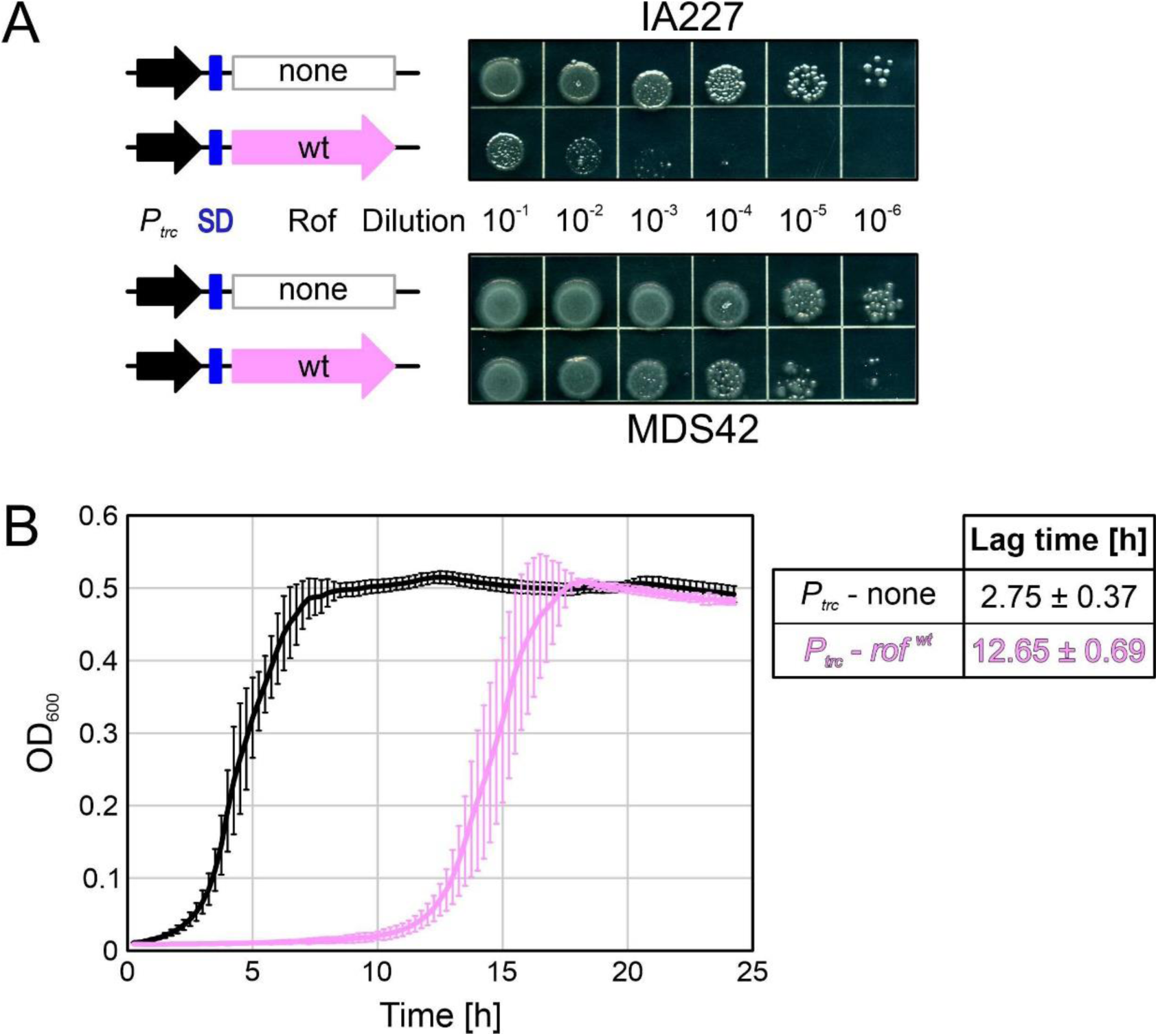
(A) A strain lacking prophages is resistant to Rof. IA227 (top) or MDS42 (bottom) cells transformed with plasmids carrying *P_trc_* promoter, with or without the *rof* gene. In these constructs, a canonical *SD* element (indicated by a blue box) is placed upstream from the ORF. Colonies from transformation plates were inoculated into LB supplemented with carbenicillin and grown for 24 h at 32 °C, 250 rpm. Serial ten-fold dilutions were plated on LB-carbenicillin and incubated at 32 °C for 24 h. A representative experiment is shown out of five biological replicates. (B) Rof expression extends the lag time but does not alter the exponential growth phase. Cells transformed with a plasmid expressing wt Rof (violet) or an empty vector (black) were grown in MOPS EZ rich defined media supplemented with carbenicillin overnight at 32 °C, diluted into fresh media supplemented with carbenicillin and 1 mM IPTG, and grown in a microplate reader at 32 °C for 24 hours. Cell growth was monitored by OD_600_ measured every 15 min. The data from four biological replicates were fit to the modified Gompertz equation (1).

**Figure S2.**
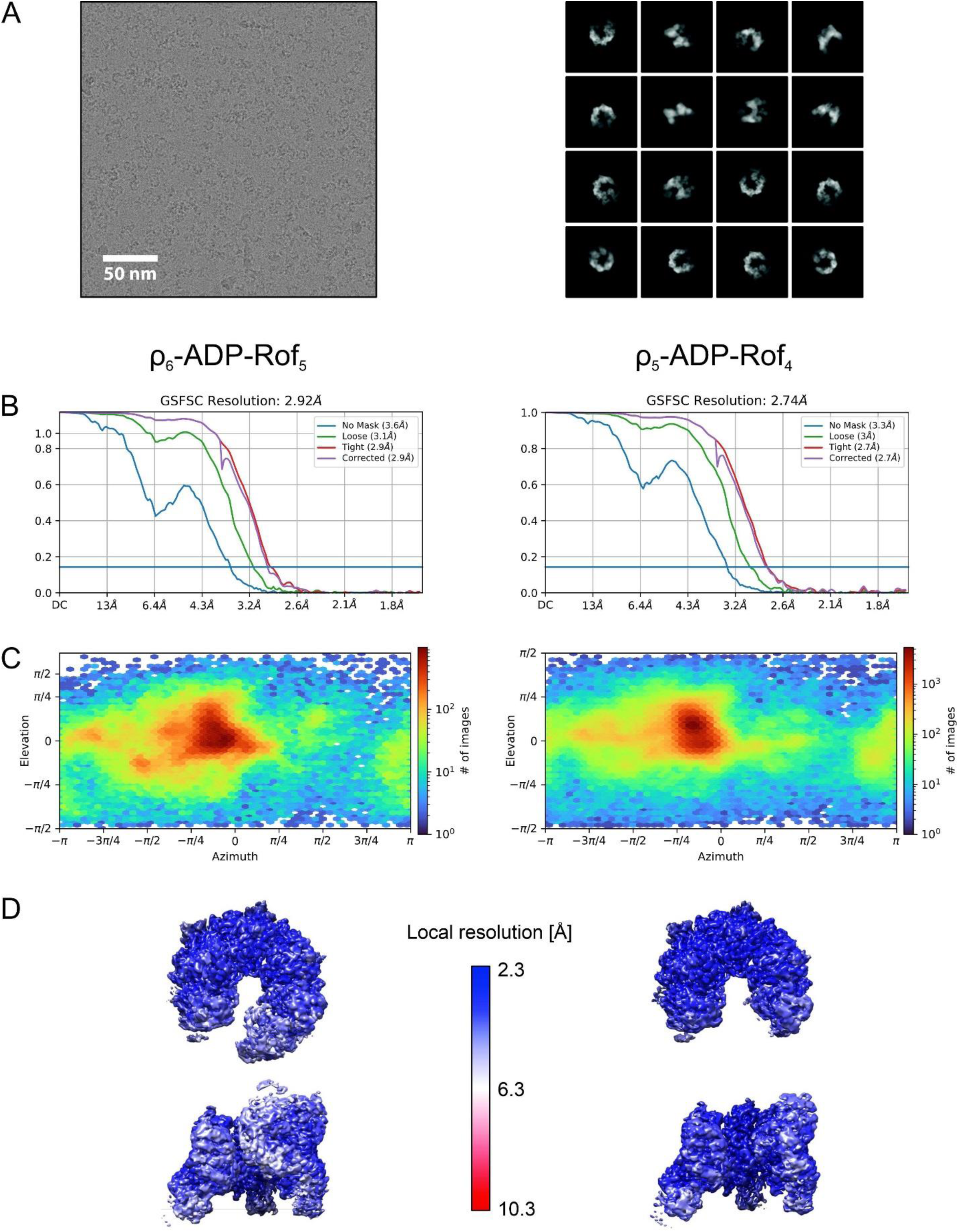
(A) Representative cryoEM micrograph (left; scale bar, 50 nm) and selected class averages (right) of ρ-ADP-Rof complexes after reference-free 2D classification. (B) Gold-standard Fourier shell correlation plots after NU refinement of the ρ_6_-ADP-Rof_5_ (left) and ρ_5_-ADP-Rof_4_ (right) reconstructions. (C) Viewing direction distributions of the ρ_6_-ADP-Rof_5_ (left) and ρ_5_-ADP-Rof_4_ (right) reconstructions. (D) Top (top) and side (bottom) views of the cryoEM reconstructions of the ρ_6_-ADP-Rof_5_ (left) and ρ_5_-ADP-Rof_4_ (right) complexes, colored according to the local resolution.

**Figure S3.**
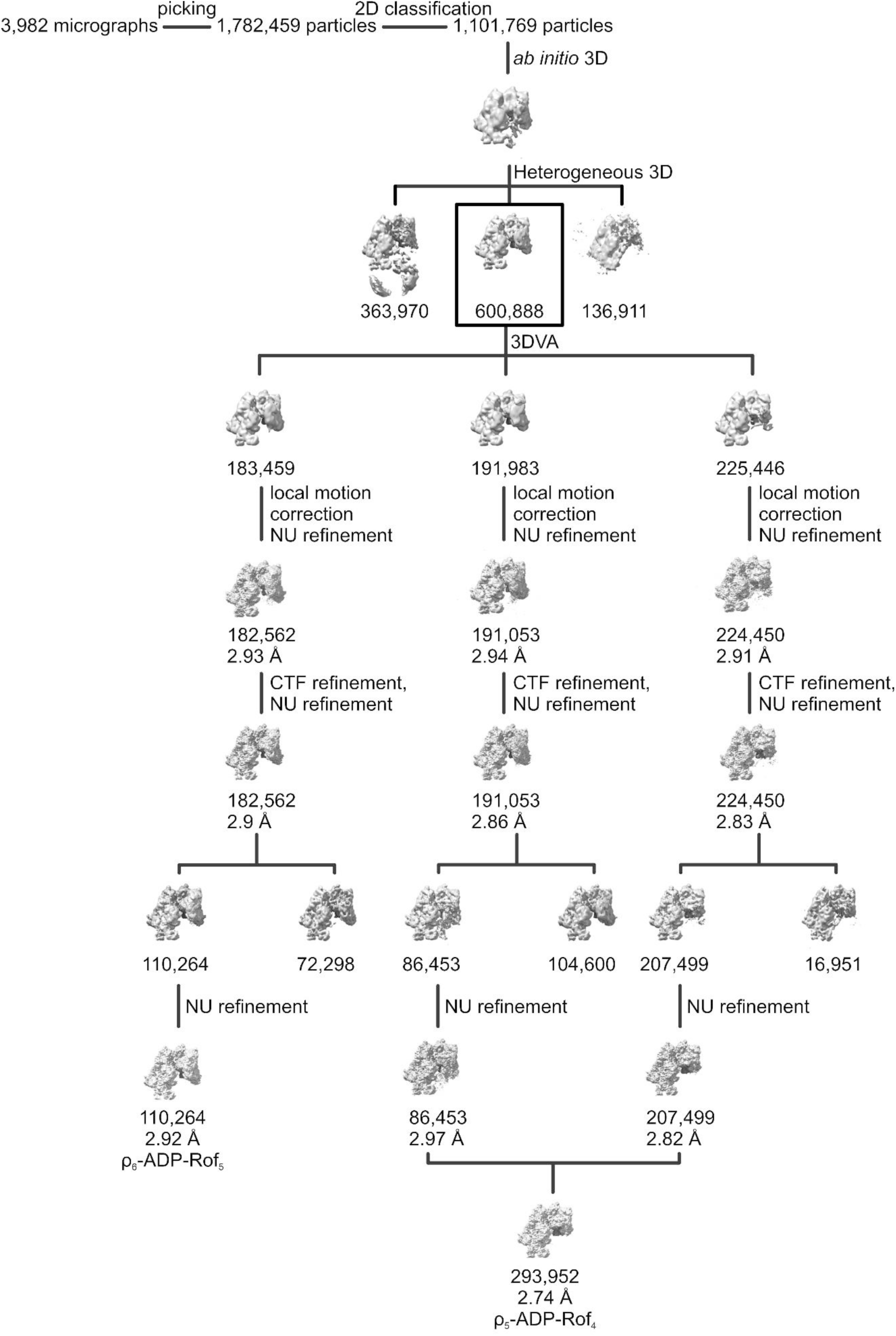
ρ-ADP-Rof cryoEM data refinement. From 3,982 micrographs a total of 1,782,459 particle images were initially picked and subjected to reference-free 2D classification. 1,101,769 particle images were selected and used for *ab initio* 3D reconstruction to generate an initial reference for heterogeneous 3D refinement. A subset of 600,888 particle images was selected for further classification by 3D variability analysis that already revealed existence of ρ_6_-ADP-Rof_5_ and ρ_5_-ADP-Rof_4_ states. Local motion correction followed by CTF refinement, heterogeneous 3D refinement, and subsequent NU refinement was applied to isolate the final particle sets and generate reconstructions of the ρ_6_-ADP-Rof_5_ and ρ_5_-ADP-Rof_4_ particles at global resolutions of 2.92 Å and 2.74 Å, respectively.

**Figure S4.**
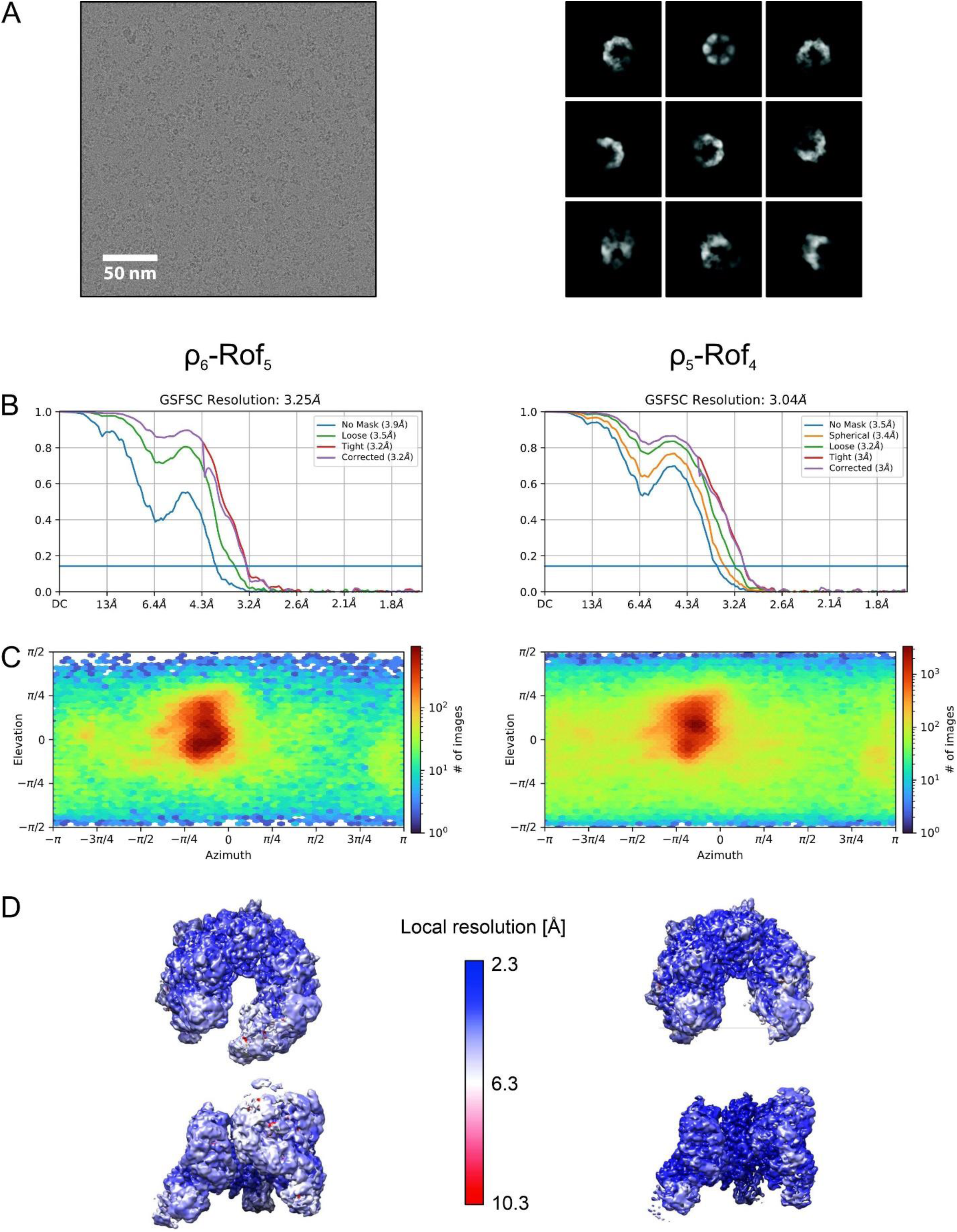
*(A)* Representative cryoEM micrograph (left; scale bar, 50 nm) and selected class averages (right) of ρ-Rof complexes after reference-free 2D classification. *(B)* Gold-standard Fourier shell correlation plots after NU refinement of the ρ_6_-Rof_5_ (left) and ρ_5_- Rof_4_ (right) reconstructions. *(C)* Viewing direction distributions of the ρ_6_-Rof_5_ (left) and ρ_5_-Rof_4_ (right) reconstructions. *(D)* Top (top) and side (bottom) views of the cryoEM reconstructions of the ρ_6_-Rof_5_ (left) and ρ_5_- Rof_4_ (right) complexes, colored according to the local resolution.

**Figure S5.**
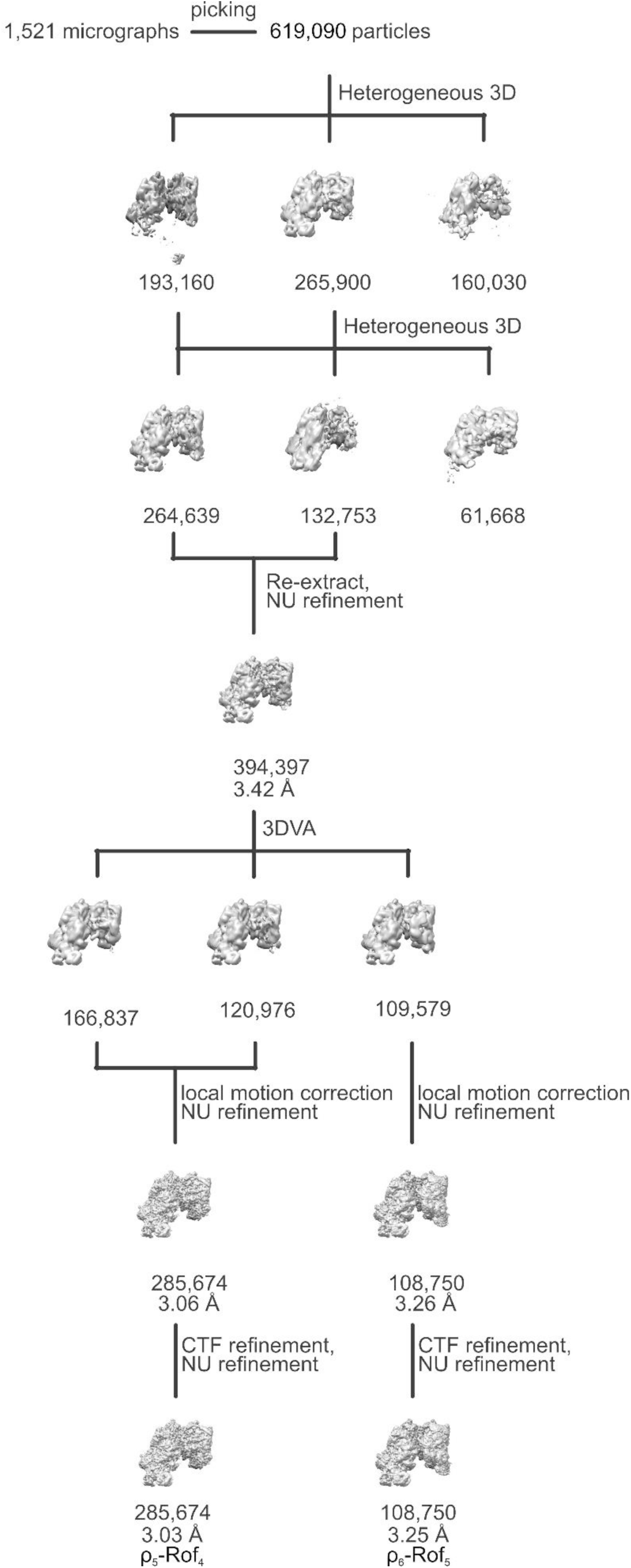
ρ-Rof cryoEM data refinement. From 1,521 micrographs a total of 619,090 particles were initially picked and subjected to iterative heterogeneous 3D refinement cycles using the ρ-ADP-Rof reconstructions as reference. 394,397 particle images were selected for re-extraction and further classification by 3D variability analysis. Local motion correction of the selected 285,674 particle images, followed by CTF refinement and NU refinement yielded a reconstruction at a global resolution of 3.03 Å.

**Figure S6.**
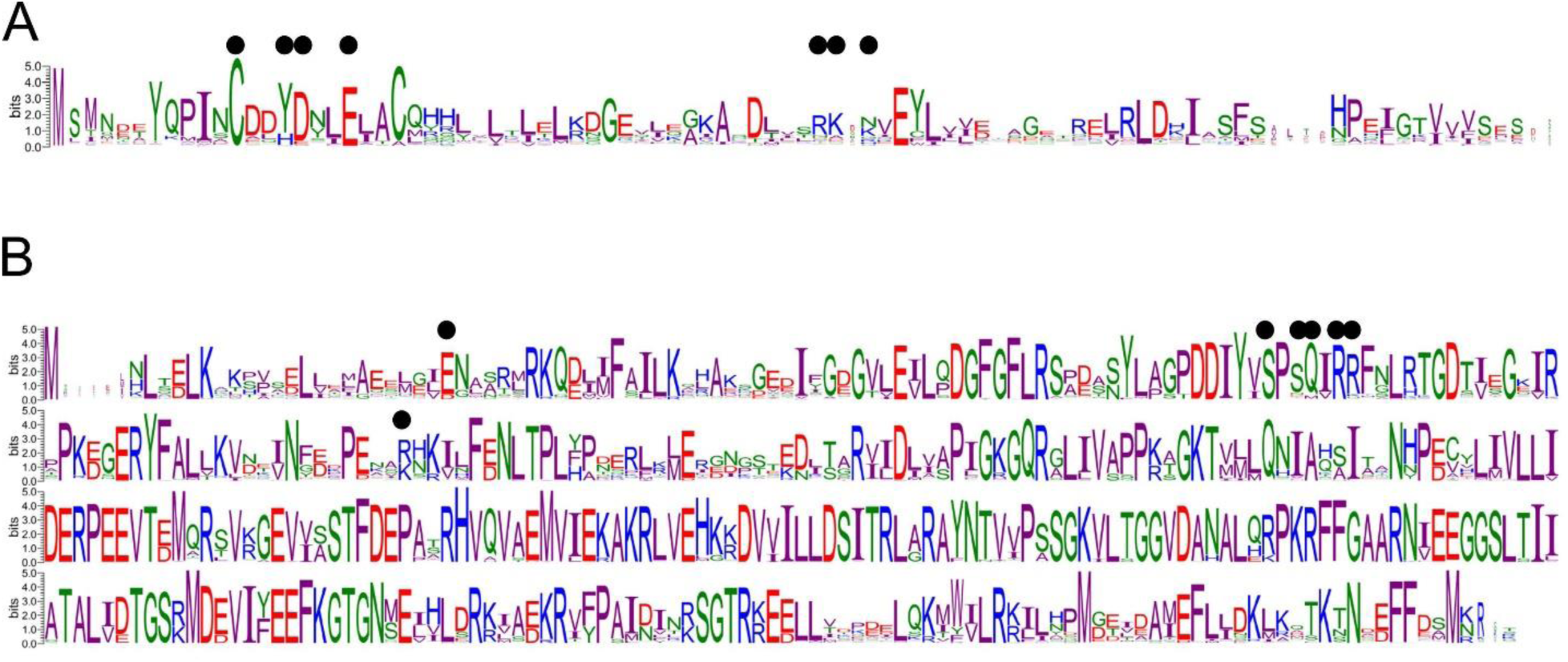
*(A)* Rof conservation. *(B)* ρ conservation. In (*A*) and (*B*), the ρ-Rof interface residues revealed by the complex structures are indicated by black dots. Sequence logos were generated by WebLogo (version 3.7.8).

**Figure S7.**
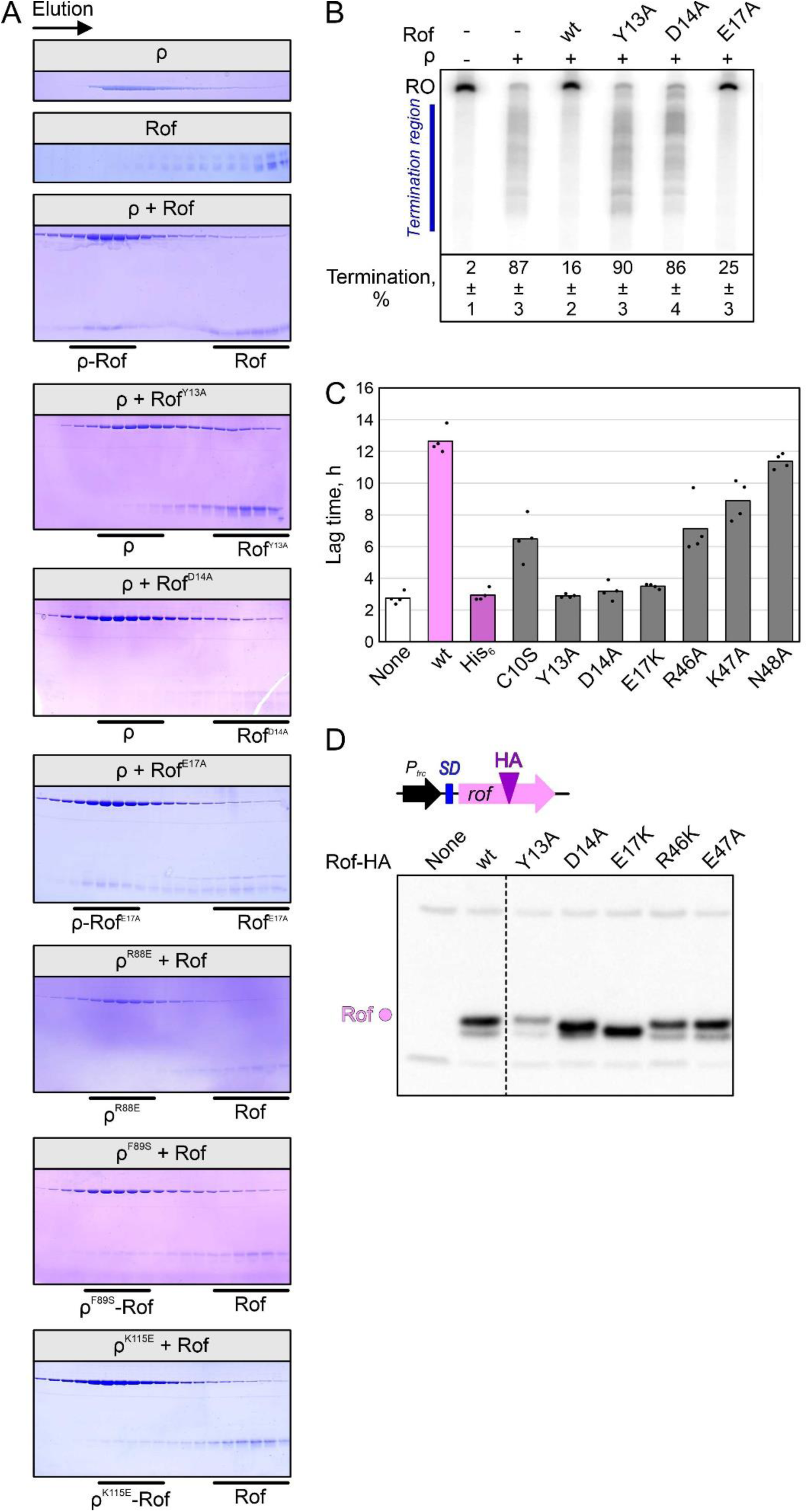
*(A)* Analytical size exclusion chromatography runs, monitoring the interaction between the indicated ρ and Rof variants. For each run, the same fractions were analyzed by SDS-PAGE. First and second panel, SEC runs of isolated ρ and Rof, respectively. Third panel, binding of ρ^wt^ to Rof^wt^. Panels 4-6, binding of indicated Rof variants to ρ^wt^. Panels 7-9, binding of indicated ρ variants to Rof^wt^. *(B)* Effects of Rof variants on ρ-dependent termination *in vitro*. Assays were done using the λ *tR1* template as in Fig. 1C. Termination efficiency values represent means ± SD of three independent experiments. *(C)* Lag times of growth for IA227 transformed with *P_trc_* plasmids expressing different Rof variants or none. The values shown are lag times calculated for four biological replicates. *(D)* Production of Rof variants containing a single HA-tag inserted after residue 57 in IA227 cells. Control experiments demonstrated that the expression of HA-tagged Rof is toxic. Rof expression following the induction with 1 mM IPTG was determined by Western blotting with anti-HA antibodies (Millipore Sigma).

**Figure S8.**
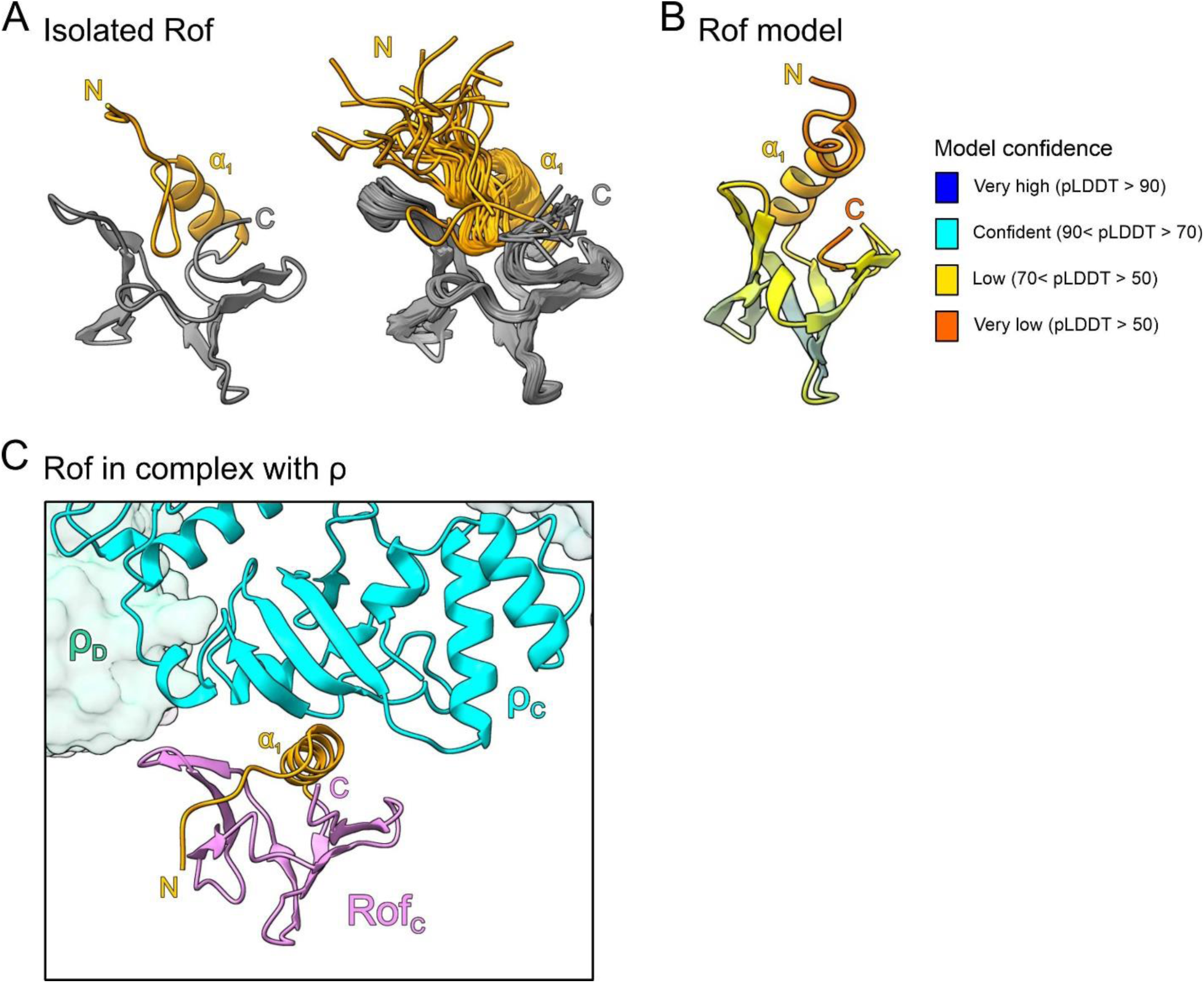
*(A)* Structure of *ec*Rof in isolation. Representative structure (left) and structural ensemble (right) of *ec*Rof determined by NMR spectroscopy (PDB ID: 1SG5) (2). The N-termini (N) and helix α_1_ are colored in orange. C, C-termini. The N-termini of isolated *ec*Rof show high flexibility and are oriented away from the protein’s core. *(B)* AlphaFold model of *ec*Rof. Rof is colored according to the AlphaFold model confidence score (pLDDT). The pLDDT score indicates structural flexibility in the N-terminal region, including helix α1. *(C)* ρ-bound *ec*Rof in the same view as isolated *ec*Rof in (*A*). The N-terminus and α_1_ of *ec*Rof bound to ρ are oriented differently compared to isolated *ec*Rof.

**Figure S9.**
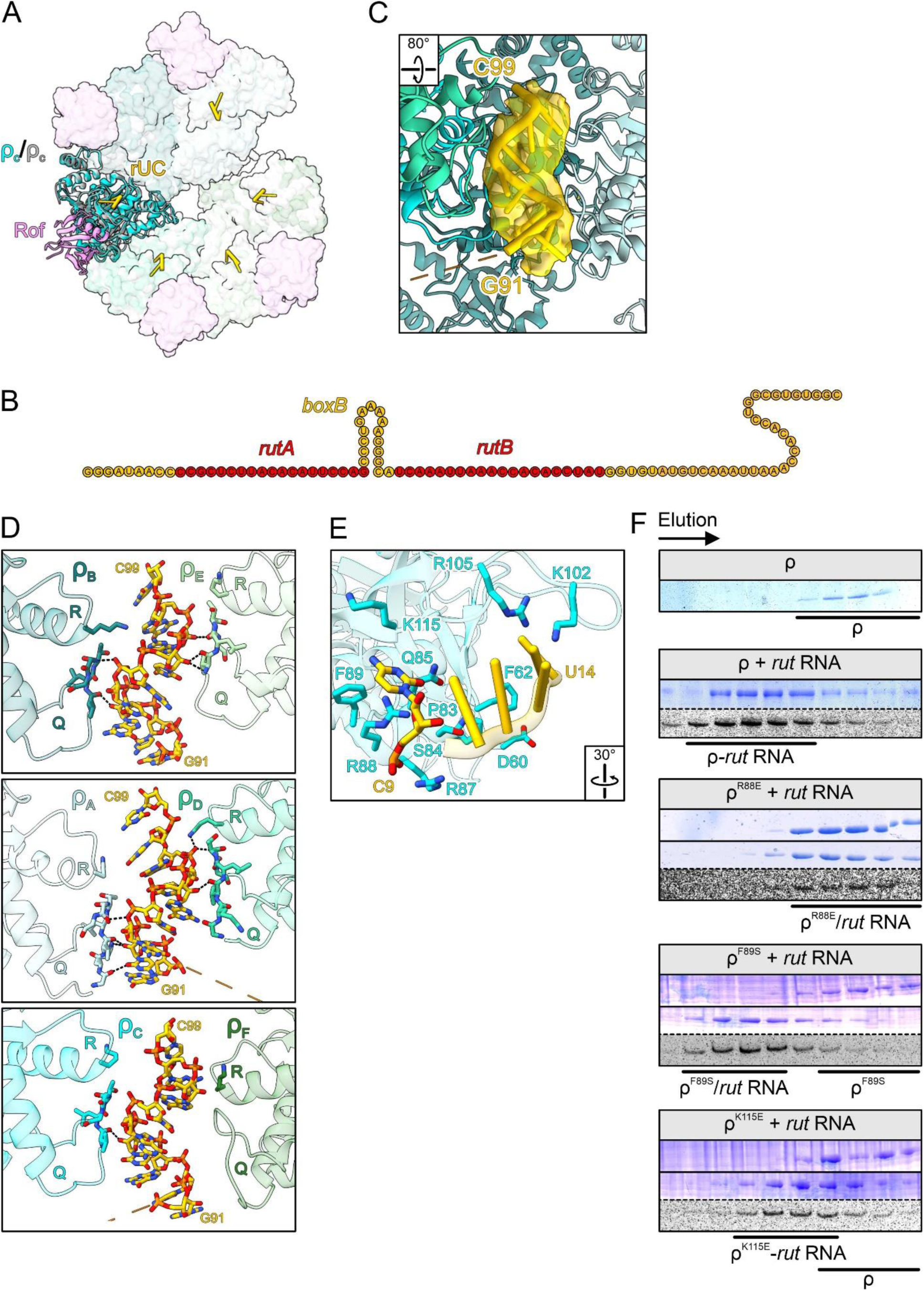
(A) Superposition of the ρ_6_-ADP-Rof_5_ structure with the structure of ρ bound to a rUC dinucleotide at the core PBS (PDB ID: 1PVO) (3). Structures are shown in cartoon (ρ_C_ and Rof_C_; ρ_C_ of rUC-bound ρ, gray) or in surface representations. Binding sites for Rof and the dinucleotide at the NTD are in close proximity but do not overlap. (B) *Rut* RNA used in this study; *rutA*, *rutB*, and the *boxB* hairpin are highlighted. (C) Experimental density for *rut* RNA at the SBS. Rotation symbols in this and in (*E*) indicate views relative to Fig. 3C. (D) ρ binds SBS RNA *via* the Q- and R-loops. RNA, Q-loop residues (281–287), and R-loop residue K326 of the ρ subunits are depicted as sticks. The RNA oriented with the 5’-end at the bottom and the 3’-end at the top. The three panels show the three pairs of ρ subunits positioned opposite to each other around the RNA. Most of the contacts are not sequence-specific and are mediated by backbone interactions of both, RNA and ρ. (E) Details of the ρ-*rut* RNA interaction along the PBS. ρ residues lining the path of the RNA are shown as sticks and labeled. Six nucleotides are bound along the OB-fold of the ρ NTD. The core PBS accommodates the dinucleotide at the 3’-end of the RNA ligand, the extended PBS accommodates the 5’-region of the RNA ligand. The 5’-terminal nt (C9) is embedded in a pocket composed of ρ residues Q85, R87, R88, F89, and K115. (F) Analytical size exclusion chromatography runs, monitoring the binding of ρ variants to *rut* RNA. For each run, the same fractions were analyzed by SDS-PAGE (protein) and 8 M urea-PAGE (RNA). First panel, SEC run of isolated ρ in the absence of RNA. Second panel, binding of ρ^wt^ to *rut* RNA. Panels 3-5, binding of the indicated ρ variants to *rut* RNA.

**Figure S10.**
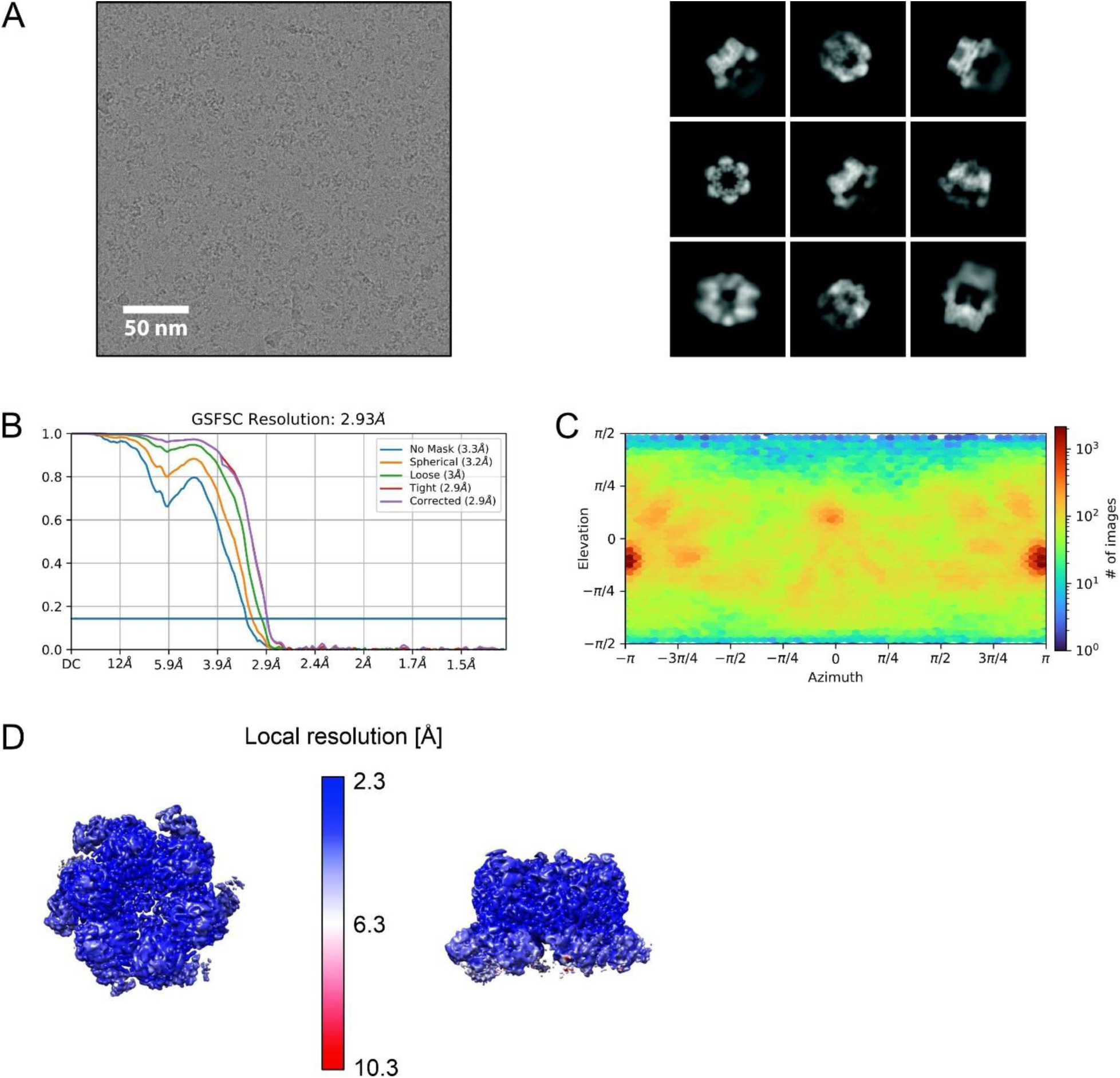
(A) Representative cryoEM micrograph (left; scale bar, 50 nm) and selected class averages (right) of ρ-*rut* RNA complexes after reference-free 2D classification. (B) Gold-standard Fourier shell correlation plot after NU refinement of the ρ-*rut* RNA reconstruction. (C) Viewing direction distribution of the ρ-*rut* RNA reconstruction. (D) Top (left) and side (right) views of the cryoEM reconstruction of the ρ-*rut* RNA complex, colored according to the local resolution.

**Figure S11.**
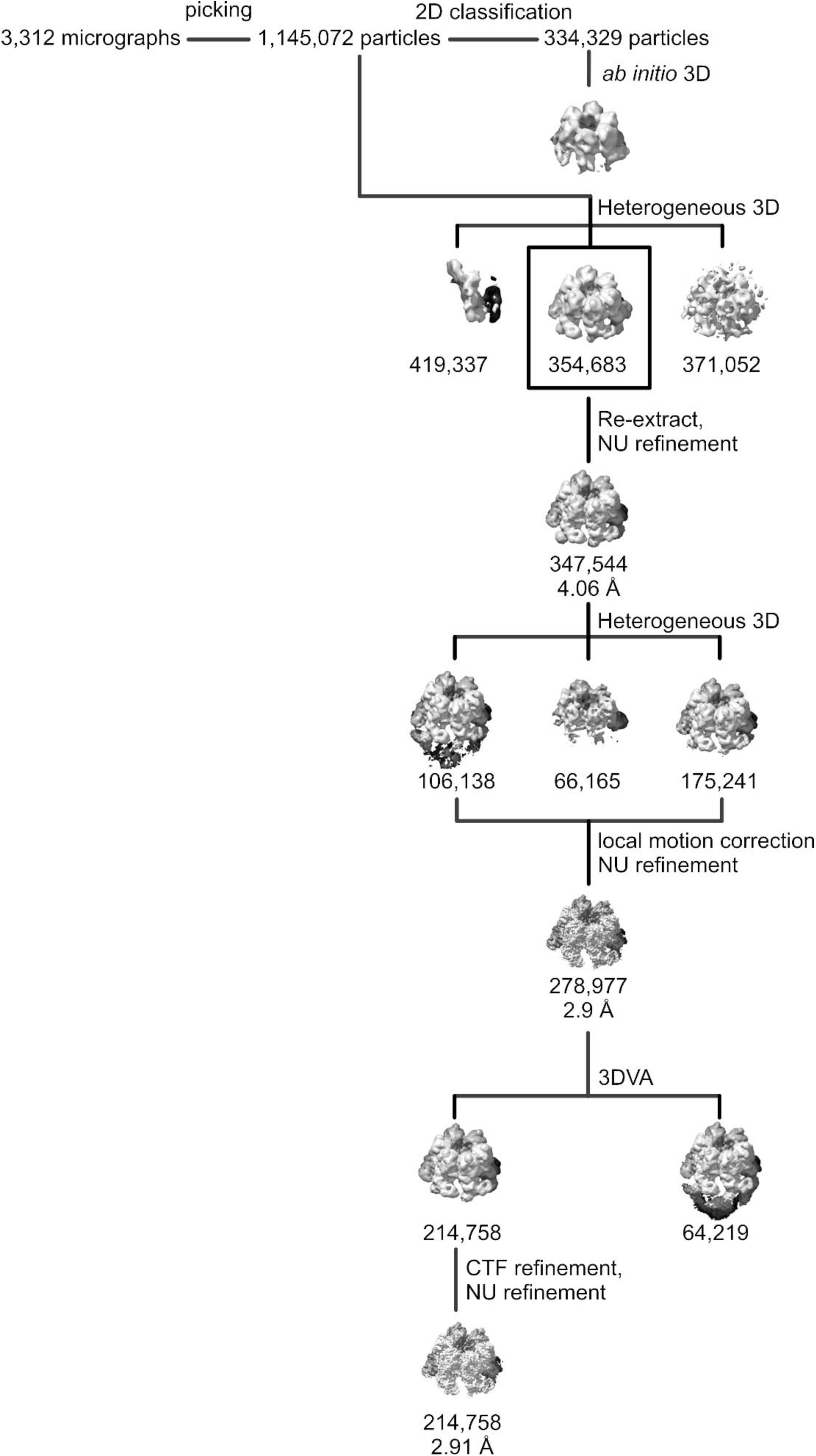
ρ-*rut* RNA cryoEM data refinement. From 3,312 micrographs a total of 1,145,072 particles were initially picked and subjected to reference-free 2D classification. 334,329 particle images were selected and used for *ab initio* 3D reconstruction to generate an initial reference for heterogeneous 3D refinement of the entire dataset. A subset of 354,683 particle images was selected for further classification by heterogeneous 3D refinement followed by local motion correction and additional classification by 3D variability analysis. CTF refinement and subsequent NU refinement of the finally selected 214,758 particle images yielded a 3D reconstruction at 2.91 Å resolution.

**Figure S12.**
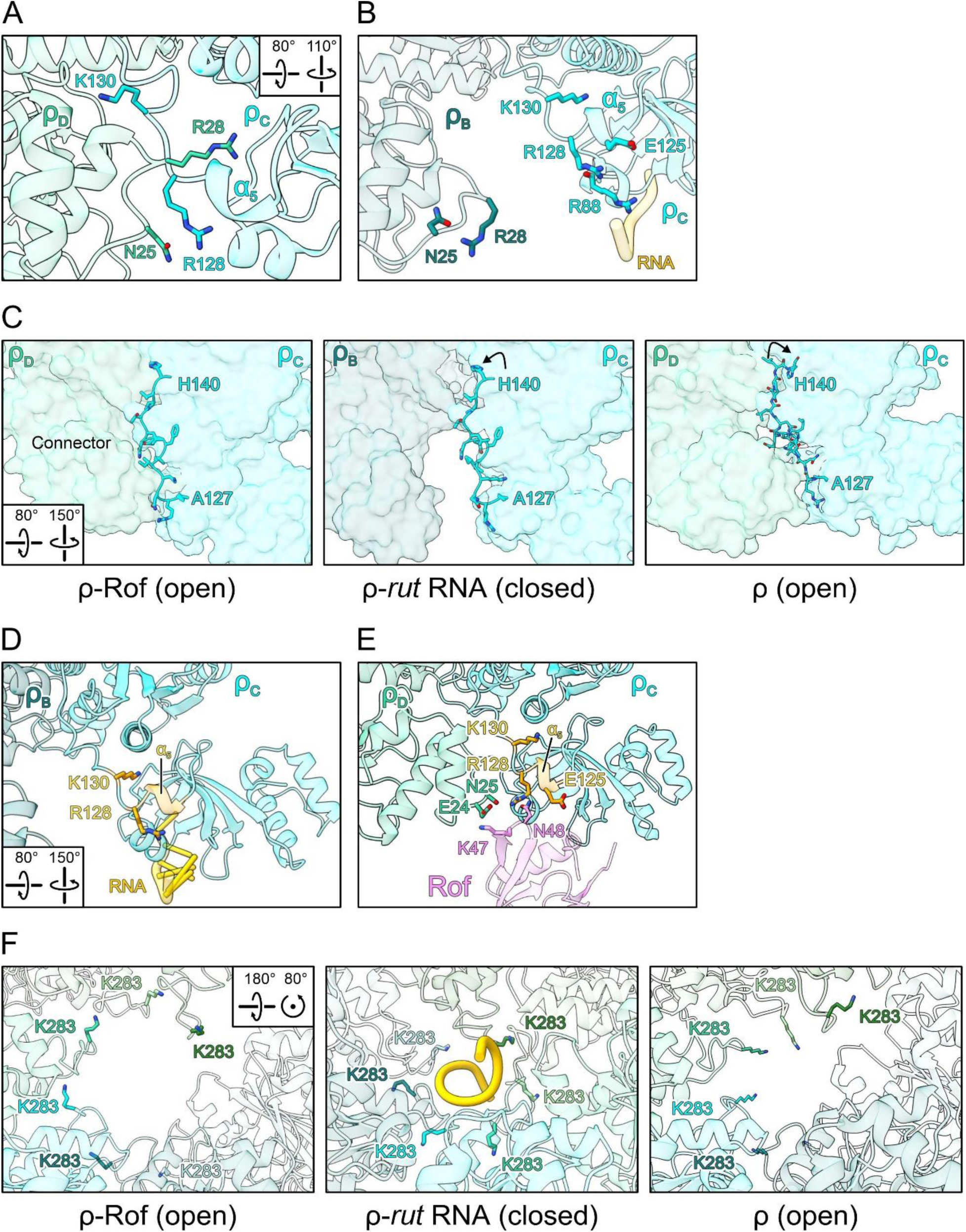
(*A, B*) Structural comparison of ρ subunit interactions in structures of open ρ bound to ATP (PDB ID: 6WA8) (4) (*A*) and closed ρ bound to *rut* RNA (*B*). In the open hexamer conformation, residue R28 from one ρ protomer (ρ_D_) is embedded in a pocket of the adjacent protomer (ρ_C_), formed partially by α_5_. In the closed hexamer conformation, R28 is ejected from that pocket and substituted by K130 of the same protomer (ρ_C_). Concomitantly, contacts between the N-terminal region of ρ_D_ and R128 of ρ_C_ are broken during ring closure (ρ_B_ an ρ_C_ in the closed hexamer). Rotation symbols indicate the view relative to Fig. 3C. (C) Comparison of the NTD-CTD connectors (sticks) in open ρ^ATP^ (left), the closed ρ-*rut* RNA complex (middle), and the open ρ_6_-ADP-Rof_5_ complex (right). Relative to open ρ^ATP^, the NTD-CTD connector (residues 127-140) in the closed ρ-*rut* RNA complex is re-aligned by one residue, most evident by the rearrangement of H140 towards the protomer interface (bent arrow). In the open ρ_6_-ADP-Rof_5_ structure, the NTD-CTD connector retains the register observed in the closed ρ-*rut* RNA complex. Rotation symbols in this and (*D*, *F*) indicate the views relative to Fig. 2B. (D) The binding of *rut* RNA to the extended ρ PBS stabilizes interactions of NTD-CTD connector within one ρ protomer, which could facilitate the disruption of ρ inter-subunit NTD-NTD contacts. (E) Rof, in contrast, fosters ρ inter-subunit NTD-NTD contacts, thereby stabilizing the open-ring configuration of the ρ NTDs. (F) Structural comparison of Q-loop conformations in open ρ^ATP^ (left), the closed ρ-*rut* RNA complex (middle), and the open ρ_6_-ADP-Rof_5_ complex (right). In open ρ^ATP^, Q-loops adopt a conformation resulting in residues K283 (sticks) pointing towards the central axis of the ρ hexamer, where they could engage in initial contacts to SBS RNA. In the closed ρ-*rut* RNA complex, K283 residues are embedded in pockets formed in part by the Q-loops of the adjacent ρ protomers. In the open ρ_6_-ADP-Rof_5_ complex, the Q-loops adopt a conformation similar to that observed in the closed ρ-*rut* RNA complex.

**Figure S13.**
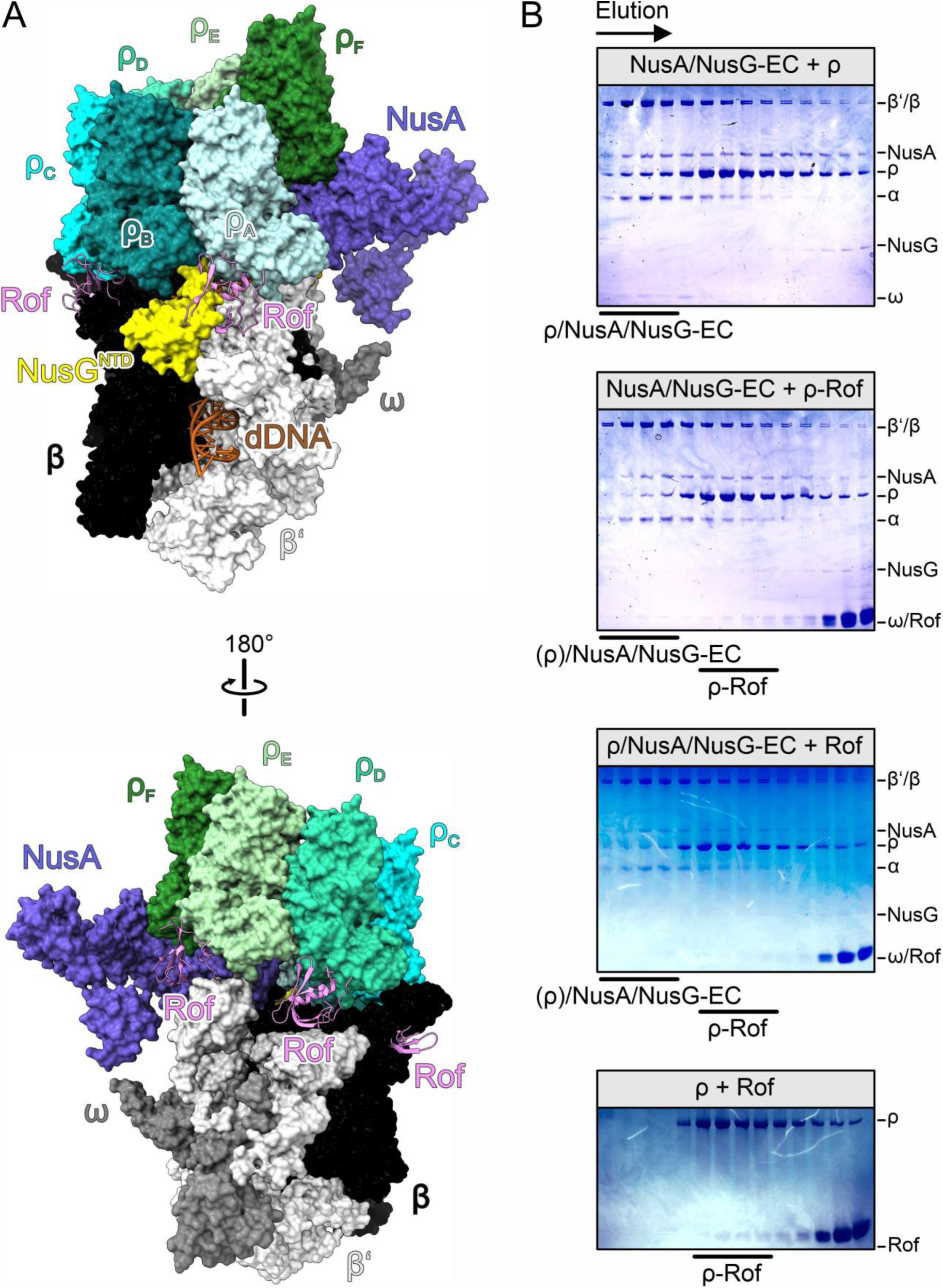
(A) Rof interferes with ρ binding to ECs. The ρ_6_-ADP-Rof_5_ structure was superimposed on the structure of a ρ/NusA/NusG-modified EC (PDB ID: 6Z9P) (5) based on the ρ_A_ protomers. RNAP subunits, different shades of grey; NusA, slate blue; NusG, yellow; downstream (d) DNA, brown. Rof sterically interferes with ρ binding to RNAP and Nus factors. (B) SDS-PAGE analysis of SEC runs, monitoring ρ binding to ECs in the absence and presence of Rof. First panel, pre-formed NusA/NusG-EC incubated with a three-fold molar excess of ρ hexamer. Second panel, pre-formed NusA/NusG-EC incubated with a three-fold molar excess (relative to ρ hexamer) of ρ-Rof complex. Third panel, pre-formed ρ/NusA/NusG-EC incubated with a ten-fold molar excess of Rof (relative to ρ hexamer). Fourth panel, ρ incubated with ten-fold molar excess of Rof (relative to ρ hexamer).

**Figure S14.**
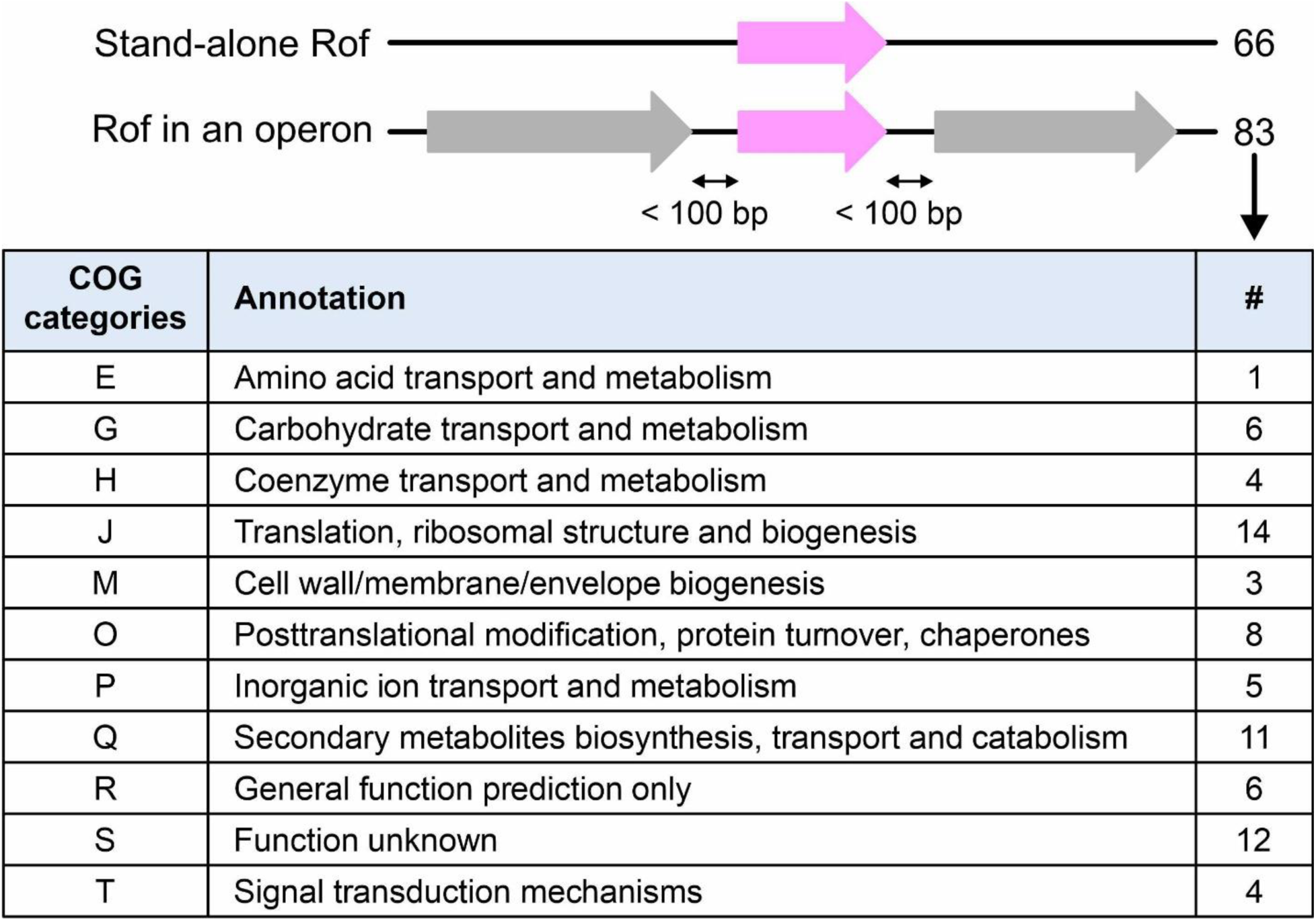
Analysis of *rof* gene neighbors collected from TREND (6) (Dataset S1*D*) shows that slightly fewer than half (66 out of 149) of the *rof* genes are not located within operons; the distance to the nearest co-directional neighbor is more than 100 bp. For those *rof* genes that are located within a hypothetical operon (83 out of 149), we assigned clusters of orthologous groups (COG) (7) to the closest neighbor of Vibrionaceae *rof,* located within 100 bp of the *rof* gene and translated from the same strand. Among 11 identified COG categories, two (J and Q), associated with the ribosome biogenesis and with the production of secondary metabolites functions, are somewhat overrepresented among the *rof* nearest neighbors.

**Table S1.** Plasmids used in this work.

| Plasmids for protein production and *in vitro* transcription |  |  |
| --- | --- | --- |
| pIA267 | $\lambda$ $P_R$ promoter – C-less A26 ITR - $\lambda$ *tR1* terminator template | (8) |
| pIA1352 | $\lambda$ $P_R$ promoter – C-less A26 ITR – Salmonella *siiE* template | This work |
| pIA586 | *E. coli* $\sigma^{70}$ expression | (9) |
| pVS10 | *E. coli* core RNAP expression; His<sub>6</sub>-tagged $\beta'$ subunit | (9) |
| pET24b-*rho* | *E. coli rho* expression | (10) |
| pETM11-*rho* | *E. coli rho* expression | (11) |
| pETM11-*rho* [R88E] | *E. coli rho* expression [R88E] | This work |
| pETM11-*rho* [F89S] | *E. coli rho* expression [F89S] | This work |
| pETM11-*rho* [K115E] | *E. coli rho* expression [K115R] | This work |
| pNS-100 | *E. coli rof* expression; N-terminal His<sub>6</sub>-TEV-tag | This work |
| pNS-100b | *E. coli rof* expression [Y13A]; N-terminal His<sub>6</sub>-TEV-tag | This work |
| pNS-100c | *E. coli rof* expression [D14A]; N-terminal His<sub>6</sub>-TEV-tag | This work |
| pNS-100d | *E. coli rof* expression [E17A]; N-terminal His<sub>6</sub>-TEV-tag | This work |
| Plasmids for *in vivo* assays; $P_{trc}$ – INSERT | | |
| pTrc99c | none | (12) |
| pIA1488 | – *E. coli rof*<sup>wt</sup> | This work |
| pIA1530 | – *E. coli rof* [Y13A] | This work |
| pIA1532 | – *E. coli* His<sub>6</sub>-*rof* | This work |
| pIA1544 | – *E. coli rof*<sup>wt</sup> [HA@57] | This work |
| pIA1545 | – *E. coli rof* [C10S] | This work |
| pIA1557 | – *E. coli rof* [Y13A;HA@57] | This work |
| pIA1574 | – *E. coli rof* [K47A] | This work |
| pIA1575 | – *E. coli rof* [R46A] | This work |
| pIA1576 | – *E. coli rof* [N48A] | This work |
| pIA1577 | – *E. coli rof* [D14A] | This work |
| pIA1589 | – *E. coli rof* [E17K] | This work |
| pIA1615 | – *E. coli rof* [D14A;HA@57] | This work |
| pIA1616 | – *E. coli rof* [E17K;HA@57] | This work |
| pIA1617 | – *E. coli rof* [R46A;HA@57] | This work |
| pIA1618 | – *E. coli rof* [K47A;HA@57] | This work |

**Table S2.**
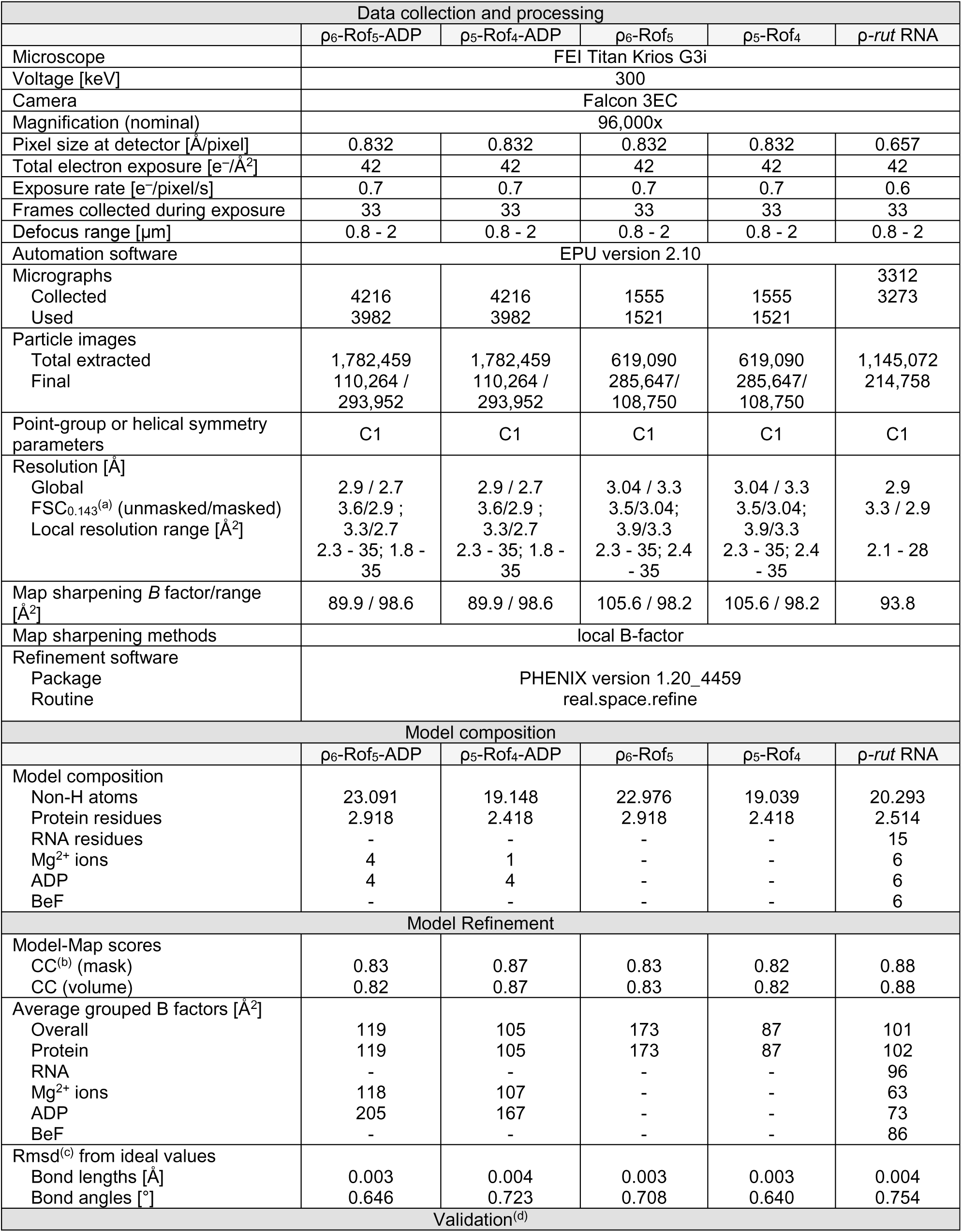
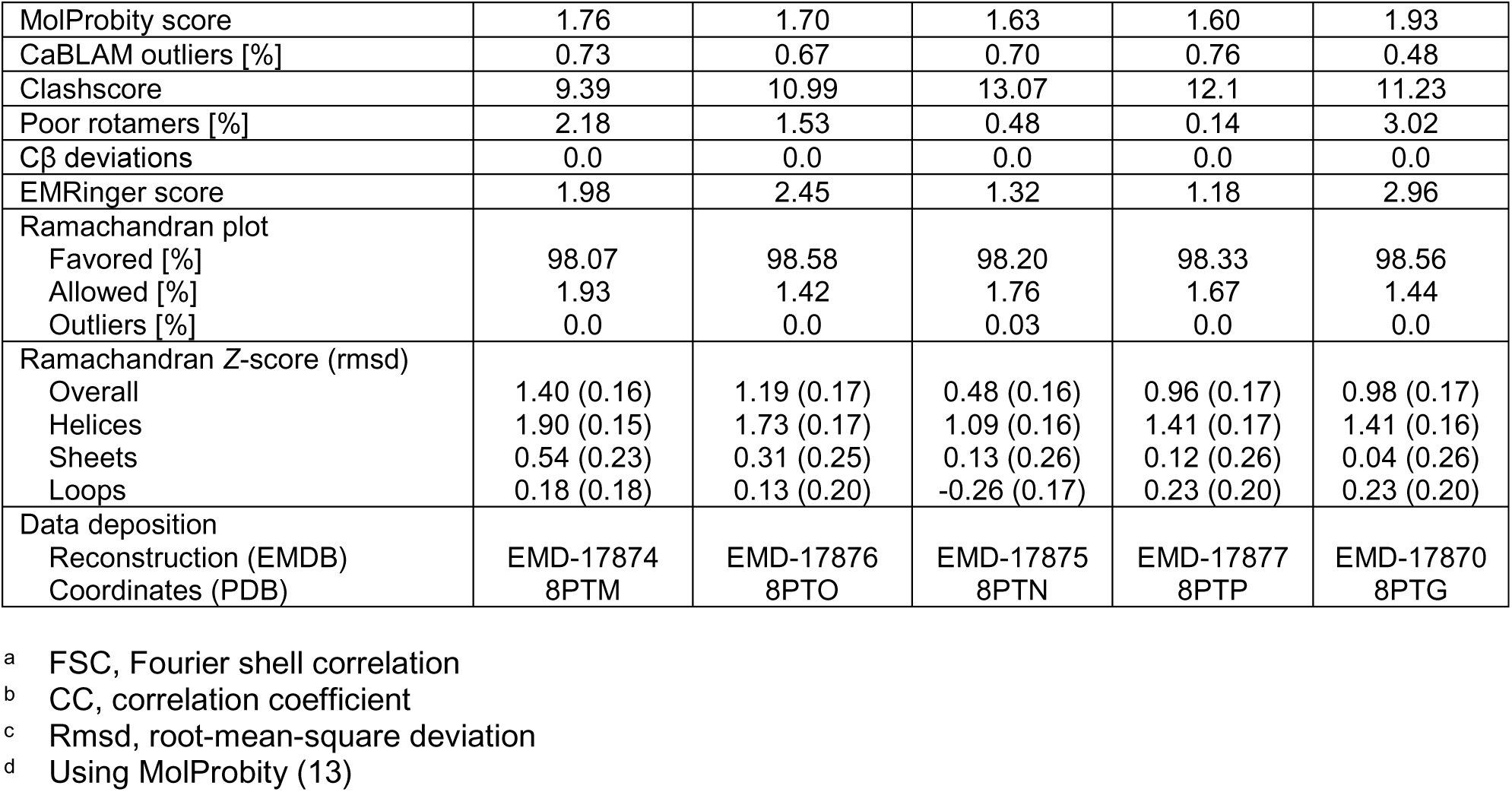
CryoEM data collection, refinement, and validation statistics.

### Dataset S1 (separate file)

Rof conservation analysis

(A) Rof distribution in families of Pseudomonadota, related to Fig. 1D. The fraction of genomes containing *rof* was estimated in Annotree.

(B) *Rof* and *yaeP* overlapping (Fig. 4E) were investigated for genomes harboring both of them in *Enterobacteriaceae*.

(C) *Rof* and *yaeP* overlapping (Fig. 4E) were investigated for genomes harboring both of them in Vibrionaceae.

(D) *Vibrionaceae rof* genomic context was collected from TREND. Note that operon ID is not continuous, which is generated by TREND. An operon is defined if *rof* has gene neighbor which is located on the same strand and within 100 bp.

(E) Rof representatives for Fig. S6*A*.

(F) ρ representatives for Fig. S6*B*.

(G) *Enterobacteriaceae* representatives for Fig. 4C.

(H) *Vibrionaceae* representatives for Fig. 4C.

## Notes

### Competing Interest Statement

The authors have declared no competing interest.

